# Deciphering the Structure and Mechanism of SaGpx: A Non-Canonical Glutathione Peroxidase from *Staphylococcus aureus*

**DOI:** 10.1101/2025.11.24.690144

**Authors:** Sushobhan Maji, Manjari Shukla, Sudipta Bhattacharyya

## Abstract

*Staphylococcus aureus* encounters massive oxidative stress during infection. To counter this, the bacterium developed robust antioxidative defense mechanism. Glutathione peroxidases (Gpx) are well characterized antioxidative enzymes in eukaryotes; however, their bacterial counterparts remain poorly explored. *S. aureus* possesses two putative Gpx genes but lacks GSH biosynthetic machinery and glutathione reductase required for canonical Gpx function, suggesting alternate electron donor system(s) may be involved. This study aimed to elucidate structure-based biochemical characterization of one of the *S. aureus* glutathione peroxidases homologs (SaGpx, Uniprot Id: Q2FYZ0) and identify its plausible electron donor system. Herein, we cloned, purified and determined the high-resolution crystal structure of SaGpx (1.5 Å resolution) using X-ray diffraction crystallography. *In vitro* biochemical characterization of the highly conserved active site amino acid point mutants, as well as their structural disposition suggests their precise roles in the enzyme’s catalysis. The crystal structure of SaGpx revealed that the enzyme adopts a canonical glutathione peroxidase fold with conserved catalytic tetrad composed of C36, Q70, W124 and N125. Also, SaGpx shows similarity with mammalian Gpx4, which was previously shown to exert phospholipid hydroperoxide peroxidase activity. Furthermore, biochemical assays suggest that SaGpx utilizes Staphylococcal thioredoxin1 as its cognate electron donor. The catalytic mechanism follows an atypical 2-cysteine peroxiredoxin-like pathway involving the formation of a sulfenic acid intermediate, followed by an intramolecular disulfide bond subsequently resolved by thioredoxin. This work provides the first structure-based biochemical characterization of a bacterial glutathione peroxidase homolog, establishing the novel structural insights of SaGpx as a noncanonical thioredoxin-dependent glutathione peroxidase.

## Introduction

Living cells continuously produces reactive oxygen species (ROS) such as hydroxyl radical (̇OH), superoxide radical 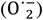, and hydrogen peroxide (H_2_O_2_) as natural byproducts of aerobic metabolism [^1^]. Multiple cellular processes generate these reactive molecules, particularly mitochondrial respiration, enzymatic reactions involved in metabolism and host immune response [^2–5^]. During bacterial infections, host immune cells increase the production of ROS. Increased level of ROS through oxidative burst, creates an oxidative stress environment intended to eliminate the invading pathogen [^5^]. Excessive ROS level becomes toxic to cells by damaging important cellular components, including proteins, lipids, and DNA, ultimately leading to cellular death [^6^]. To survive in this hostile environment, organisms developed antioxidative defense mechanism. This antioxidative defense mechanism is composed of an array of enzymes including catalase, superoxide dismutase, and peroxidases [^7–9^]. Among these antioxidants, thiol dependent peroxidases play a significant role in maintaining cellular redox homeostasis by neutralizing hydroperoxides (ROOH). The thiol peroxidase family consists of two major enzyme groups namely glutathione peroxidase (Gpx) and peroxiredoxin (Prx) [^10^]. These enzymes are the front runner proteins of specific enzyme cascade (Scheme 1) and reduce hydrogen peroxides and organic hydroperoxides into water and respective non-toxic alcohols, using evolutionarily conserved cysteine residues that undergoes cycles of oxidation and reduction [^11–13^]. Both these families of enzymes take electron from the ultimate source NADPH, generated during the metabolism of glucose via the pentose phosphate pathway [^14^]. However, the electron relay from NADPH to the terminal Gpx/Prx is mediated by different reducing agents. Glutathione-dependent Gpx utilizes glutathione reductase (GR) and glutathione (GSH), whereas thioredoxin-dependent Gpx utilizes thioredoxin reductase (TR) and thioredoxin (Trx) in a similar mechanism as of 2-cys Prx [^12^].

**Scheme 1.**
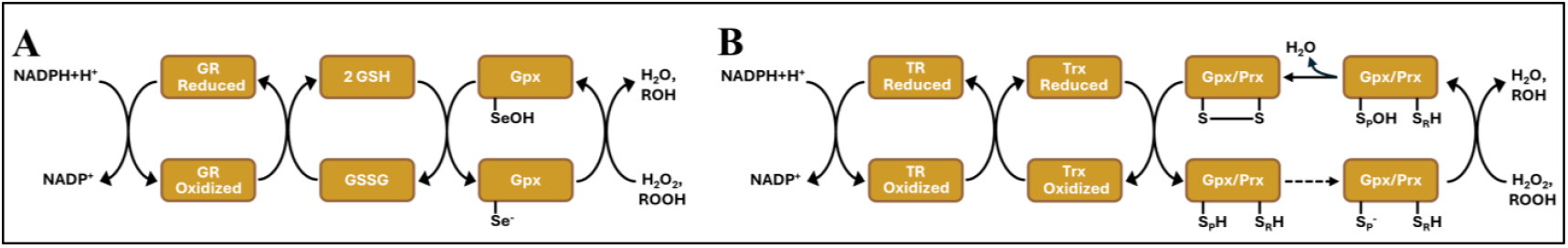
Catalytic mechanism of Thiol Peroxidase (Gpx/Prx). The electron flows from NADPH+H^+^ to ultimately peroxides converting them into respective alcohols/water. Electrons from NADPH+H^+^ takes two routes to neutralize substrate hydroperoxides: (1A) GR→GSH→Gpx in case of GSH-dependent Gpxs, and (1B) TR→Trx→Prx in case of and 2-Cys Prx and Trx-dependent Gpxs (TR→Trx→Gpx). GR: glutathione reductase; GSH: reduced glutathione; GSSG: oxidized glutathione; Gpx: glutathione peroxidase; TR: thioredoxin reductase; Trx: thioredoxin; Prx: peroxiredoxin.

Gpx is well characterized both in mammals as well as in other eukaryotes. In human, eight copies of Gpx are present. Due to their widespread localization in the body and antioxidant properties, human Gpx’s takes part in different important structural role, including the protection of reproductive and sensory functions amongst many [^11,13^]. Despite extensive characterization of eukaryotic Gpx enzymes, their bacterial counterparts remain elusive and poorly understood due to the lack of experimentally determined structural information. Studies on *Streptococcus pyrogens* shows Gpx plays an important role in bacterial pathogenesis [^15^]. In contrast, recent studies on *Listeria monocytogenes* shows a counter-intuitive effect of Gpx on the antioxidant capacity and pathogenicity. However, it has also been shown that, Gpx from *Listeria monocytogenes* has regulatory role on virulence factors of the pathogen [^16^]. Nevertheless, these findings collectively suggest that bacterial Gpx enzymes may play important roles in pathogenesis during host-pathogen interactions.

*Staphylococcus aureus*, a clinically significant pathogen, encounters intense oxidative stress during infection and relies on antioxidant enzymes for survival in hostile intracellular microenvironment of the host [^17,18^]. Amongst several other antioxidant systems, the expression of glutathione peroxidase in *S. aureus* is also upregulated immediately after phagocytosis [^19^]. Two copies of putative Gpx genes have been annotated in the *S. aureus* genome [^20^]. However, their biochemical roles remain unexplored. Intriguingly, *S. aureus* lacks the classical glutathione-based electron donor system (as depicted in Scheme 1A) typically associated with Gpx function. The bacterium does not possess glutathione biosynthetic machinery [^21^]. Recent work by Lensmire et. al. 2023 [^22^], identified a glutathione importer in *S. aureus*. This importer is associated with a γ–glutamyl transpeptidase, which is reported to provide nutrient sulfur requirement of the organism by cleaving imported GSH/GSSG. However, *S. aureus* lacks the immediate reducing agent of the oxidized glutathione GSSG, the enzyme, glutathione reductase [^21^]. The absence of both endogenous glutathione synthesis and glutathione reductase enzyme makes the recycling of Gpx through the canonical GR→GSH→Gpx cascade unfavourable for hydroperoxide stress mitigation in *S. aureus*. These observations suggest that *S. aureus* Gpx homologs must utilize alternative electron donors for their catalytic regeneration, in the context of the critical role of Gpx homologs in bacterial pathogenesis [^15^]. This hypothesis is supported by the studies of Gpx homologs in other organisms like poplar [^23^], *Saccharomyces cerevisiae* [^24^] and *Drosophila melanogaster* [^25^] among many, where Gpx can use thioredoxin instead of glutathione for recycling and continuation of the enzyme cascade. These thioredoxin-dependent Gpx enzymes employs a catalytic mechanism similar to atypical 2-cys peroxiredoxin [^26^] which includes three key steps: 1) nucleophilic attack by the peroxidative cysteine (C_P_) on the polarized peroxyl bond of the substrate hydroperoxides. This attack leads to the release of alcohol (R-OH) or water with subsequent formation of enzyme-sulfenic acid intermediate (-S-OH); 2) resolution of the enzyme-sulfenic acid by the resolving cysteine (C_R_) leading to formation of an intramolecular disulfide bond; and 3) reduction of this intramolecular disulfide bond by thioredoxin to regenerate the active enzyme. Given the important role of Gpx in bacterial pathogenesis and virulence, as well as the absence of cognate electron donor of Gpx in *S*. aureus, _motivates us to explore structure-based functional attributes of Gpx homologs in *S. aureus*.

Among the two putative Gpx genes in *S. aureus* strain NCTC8325, we focused on SAOUHSC_01282, which contain three cysteine residues rather than SAOUHSC_02949, which possesses only one cysteine. The three-cysteine variant more closely resembles to glutathione-independent thiol peroxidases found in other organisms. We named this Gpx as SaGpx. Prior to this study, no previous high-resolution structure of a bacterial Gpx homolog has been reported. In this study, we determined the high-resolution crystal structure of SaGpx (at 1.5 Å resolution) using x-ray diffraction crystallography. Combined with site-directed point mutagenesis of the highly conserved active site amino acid residues and corresponding biochemical characterization, our structural analysis reveals the functional mechanism of this enzyme, representing the first structure based functional characterization of a bacterial glutathione peroxidase.

## Result

### 1. Multiple sequence alignment analysis reveals conserved glutathione peroxidase like features present in Staphylococcal glutathione peroxidase homolog (SaGpx, Uniprot Id: Q2FYZ0)

A multiple sequence alignment was carried out for the putative glutathione peroxidase (SaGpx) from *Staphylococcus aureus* NCTC8325, using functionally and structurally characterized Gpx homologs as reference sequences. The sequence alignment data shows SaGpx harbours similar characteristic features as of other Gpx homologs. Notably, SaGpx shows more than 40% overall sequence similarity with glutathione peroxidases or glutathione peroxidase-like thiol peroxidases from *Saccharomyces cerevisiae* (PDB ID 3CMI), *Populus trichocarpa x Populus deltoides* (PDB ID 2P5Q), *Trichoderma reesei* (PDB ID 6VPD), *Trypanosoma cruzi* (PDB ID 3E0U), *Schistosoma mansoni* (PDB ID 2WGR), *Trypanosoma brucei* (PDB ID 2VUP) and human Gpx-4 (PDB ID 2OBI). Prior studies on these Gpx homologs have established the importance of a conserved catalytic tetrad comprising cysteine (C), glutamine (Q), tryptophan (W), and asparagine (N) residues [^23,24,27–31^]. In SaGpx, amino acid residues C36, Q70, W124, and N125 align with equivalent residues in known Gpx homologs **(Fig. 1)**, establishing their conservation all throughout, suggesting their potential role in enzyme catalysis. Sequence alignment data also highlights the presence of three conserved motifs throughout the Gpx homologs as reported in Redoxibase database [^32^]. These conserved segments, referred to as Gpx signature motifs are 1. (G[K/R]x[L/V][I/L]I[V/E/T]NVA[S/T/A][E/Q/L/Y][C/U]G[L/T]T), 2. (LAFPCNQF), and 3. (WNF(S/T)KF). These conserved motifs possess some of the active site residues and are crucial for the catalytic activity of Gpx. These motifs, along with the tetrad, are located within conserved structural elements that is active-site loops known to be essential for glutathione peroxidase function.

**Figure 1.**
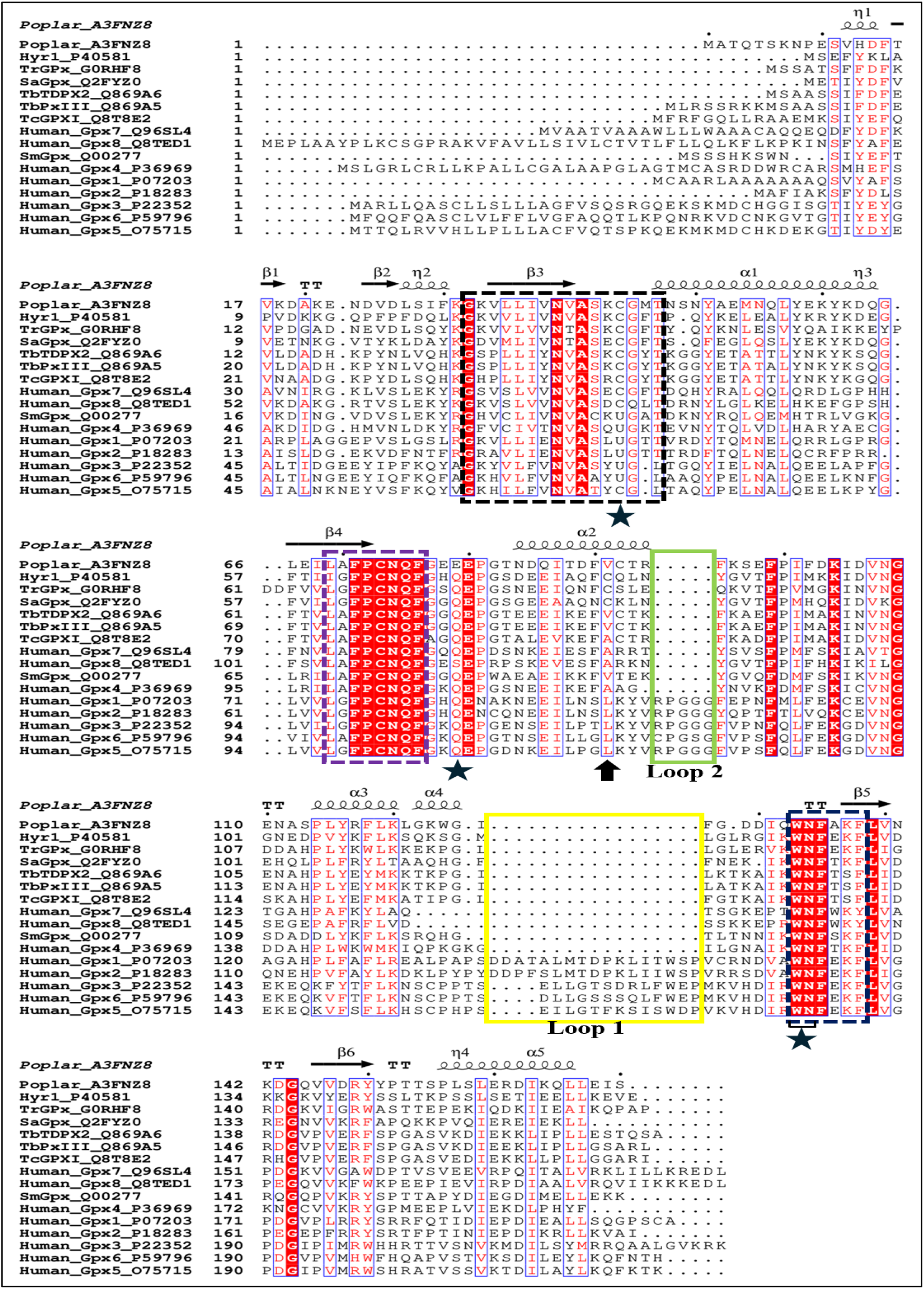
Multiple sequence alignment based on the amino acid sequence of SaGpx with homologs (with known experimentally determined structures) from different organisms. Protein sequences and corresponding crystal/NMR structures are accessible in the Uniprot database under the following accession number: *Populus trichocarpa x Populus deltoides* (A3FNZ8, PDB ID 2P5Q), *Saccharomyces cerevisiae* (P40581, PDB ID 3CMI), *Trichoderma reesei* (G0RHF8, PDB ID 6VPD), *Trypanosoma brucei* (Q869A6, PDB ID 2VUP), *Trypanosoma brucei* (Q869A5, PDB ID 2RM5), *Trypanosoma cruzi* (Q8T8E2, PDB ID 3E0U), *Schistosoma mansoni* (Q00277, PDB ID 2WGR), and human Gpx-4 (P36969, PDB ID 2OBI). Gpx homologs were selected based on the amino acid sequence similarity (more than overall 40% sequence identity with SaGpx amino acid sequence) and already having crystal/NMR structure. Apart from that, human Gpx homologs, Gpx1 (P07203, PDB ID 2F8A), Gpx2 (P18283, PDB ID 2HE3), Gpx3 (P22352, PDB ID 2R37), Gpx5 (O75715, PDB ID 2I3Y), Gpx6 (P59796), Gpx7 (Q96SL4, PDB ID 2P31), and Gpx8 (Q8TED1, PDB ID 3CYN) were also considered for sequence alignment for comparative analysis. Dotted squared boxes represents the Gpx signature motifs; black dotted square, motif 1 (G[K/R]x[L/V][I/L]I[V/E/T]NVA[S/T/A][E/Q/L/Y][**C/U**]G[L/T]T), purple dotted square, motif 2 (LAFPCNQF), and dark blue dotted square, motif 3 (WNF(S/T)KF). Solid Yellow box represents loop1 while green box represents loop 2. (★) represents the catalytic tetrad residues composed of C36, Q70, W124 and N125 respectively. (↑) represents the nonaligned resolving cysteine, C82. The sequence alignment was mapped with known secondary structural information of poplar Gpx. The alignment was performed with ClustalW and analysed with Espript.

### 2. Cloning, over-expression and purification of Wild type SaGpx, its mutants (C36S SaGpx, C64S SaGpx, C82S SaGpx, N70A SaGpx, W124A SaGpx, N125A SaGpx, (C64S+C82S) SaGpx), and plausible electron doners (SaTrx1, SaTrx2 and SaTR)

The presence of Gpx hallmark features in SaGpx amino acid sequence motivates us to clone, over-express and purify SaGpx for its structural and functional characterization. Along with SaGpx, other possible electron donor proteins of the staphylococcal thiol-based antioxidative defense pathway for example, Staphylococcal Thioredoxin 1 (SaTrx1), Staphylococcal Thioredoxin 2 (SaTrx2) and Staphylococcal Thioredoxin Reductase 1 (SaTR) were also cloned, overexpressed and purified. SaGpx contains three cysteine residues in its amino acid sequence. To assess their contribution in catalysis, site-directed mutants were also generated in which each cysteine residue was individually substituted with serine, yielding C36S, C64S, and C82S point mutants, as well as the double mutant where both of the plausible resolving cysteines were mutated (C64S+C82S) was also generated. In addition, to dissect the functional importance of amino acid residues forming the putative catalytic tetrad of SaGpx, alanine substitution mutants were constructed for Q70, W124, and N125, resulting in N70A, W124A, and N125A point mutants of SaGpx. The ORFs of all these proteins were cloned in a suitable *E. coli* expression vector. Upon successful cloning, these proteins were over-expressed under IPTG induction (100µM) at 37º C and harvested after post-induction incubation of 4 hours at the same temperature. The harvested *E. coli* cells with over-expressed target proteins were then sonicated and the soluble protein fraction was collected upon high-speed centrifugation (22,300 g for 1 hour). The soluble protein fractions were then purified by passing through the Ni-NTA based IMAC column at different step gradients of imidazole concentrations. The Ni-NTA purified proteins were further subjected to size exclusion chromatography (SEC) to obtain homogeneously purified protein samples. The purity of desired protein constructs was ensured by SDS-PAGE (12% or 15%) analysis. The in-solution homogeneous fractions of the purified proteins were collected based on sharp and single SEC elution peak at λ_Max_ = 280nm **(Fig. S1)**. The SEC purified proteins were snap-frozen by liquid nitrogen and stored at −80°C until their use for further functional characterization. However, freshly purified proteins were used for crystallization purposes.

### 3. Wild type SaGpx shows thiol dependent peroxidase activity, while the mutants exhibit altered catalytic activity

Sequence alignment analysis classified SaGpx as a member of the thiol peroxidase enzyme family. To experimentally validate this, the thiol peroxidase activity of recombinant wild type SaGpx was assessed. If SaGpx possesses peroxidase activity, then catalytically active SaGpx (the fully reduced form of the enzyme) should break down substrate hydroperoxides into their respective alcohol (or water in case of hydrogen peroxide) while simultaneously undergoing self-oxidation. In doing so, the enzyme itself got converted into disulfide-linked oxidized state, the inactive form of SaGpx. Since, we are characterizing SaGpx as a novel bacterial glutathione peroxidase, we hypothesized that the enzyme would require a reducing agent to regenerate its active form (the fully reduced form of the enzyme), similar to other members of the Gpx family. To test this hypothesis, initially, the SaGpx enzyme activity assay was performed in the absence of any exogenous reducing agent. In the absence of any exogenous reducing agent, it is expected that the oxidized state of SaGpx could not be recycled back into its catalytically active fully reduced state. Under this experimental condition, if SaGpx exhibits peroxidase activity, the extent of its self-conversion from the fully reduced active state to catalytically inactive oxidized state should be directly proportional to the amount of substrate hydroperoxide catalytically converted to alcohol and/or water. This conversion from reduced to oxidized state of SaGpx was monitored using Ellman’s reagent or DTNB which is a reagent used for quantifying free sulfhydryl (-SH) groups in solution. DTNB reacts with the conjugate base (R-S^−^) of the free sulfhydryl/thiol group, releasing a yellow-coloured species called TNB (2-nitro-5-thiobenzoic acid), which could be spectrophotometrically detected by its characteristic absorption at 412 nm. With the increasing substrate hydroperoxide concentrations, the conversion rate of the catalytically inactive oxidized form of the enzyme would be higher and lesser number of free thiol group would be available to interact with DTNB, leading to less TNB generation. This would provide direct evidence of SaGpx’s ability to undergo the characteristic oxidative catalytic cycle which is expected from a functional thiol peroxidase enzyme. The decreasing intensity in the absorbance at 412nm **(Fig. 2A)** with increasing tertiary-butyl hydroperoxide (t-BOOH) concentrations supports the decrease in free thiol (-SH) group as a result of disulfide bond formation, which in turn supports the catalytic transition of SaGpx from reduced state to oxidized state after the catalytic conversion of t-BOOH to t-BOH (tertiary-butyl alcohol) and water. This DTNB based thiol peroxidase activity assay confirmed that SaGpx actively catalyses the reduction of hydroperoxide substrates, thereby establishing its functional classification as a thiol peroxidase enzyme.

**Figure 2.**
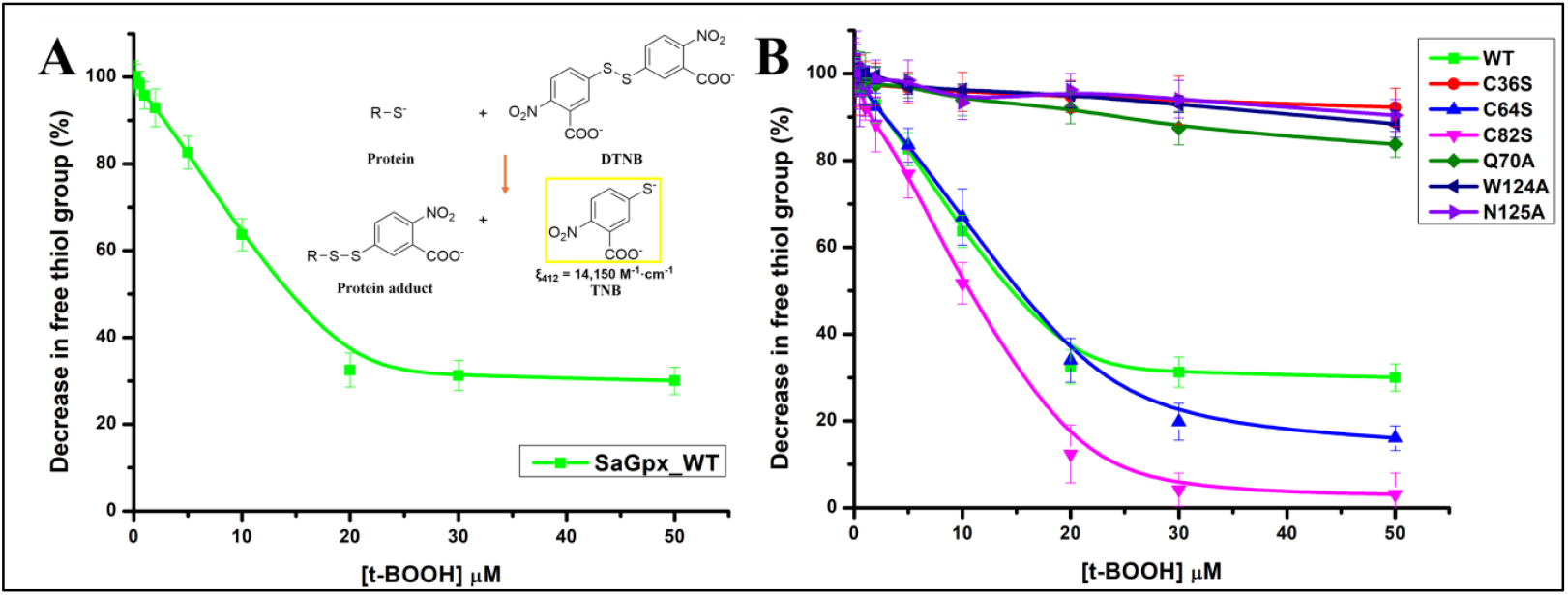
Assessment of thiol dependent peroxidase activity of SaGpx and its point mutants. (2A) Wild type SaGpx (SaGpx_WT) shows peroxidase activity in the presence of substrate tertiary-butyl hydroperoxide (t-BOOH). Upon increasing substrate concentration, SaGpx_WT shows a decrease in absorbance at λ_Max_ = 412 nm upon DTNB addition, indicating oxidation of free thiol as a function of increasing t-BOOH concentrations. (2B) Comparison of thiol-dependent peroxidase activity of different active site point mutants of SaGpx along with the wild type of enzyme. C36S (demonstrated by red traced line) mutant shows no such change in free thiol group compared to wild type (demonstrated by green traced line). C64S (demonstrated by blue traced line) shows decrease in free thiol content similar to wild type, however C82S (magenta) shows more decrease in free thiol content (see section 3 of result section for more details). In contrast, Q70A (demonstrated by olive traced line), W124A (demonstrated by navy blue traced line), and N125A (demonstrated by violet traced line) mutants show no prominent change in free thiol content upon increasing substrate concentration.

To further explore the functional relevance of the highly conserved amino acid residues involved in the catalytic activity of SaGpx, the same peroxidase activity assay was performed with SaGpx mutants: C36S, C64S, C82S, Q70A, W124A, and N125A **(Fig. 2B)**. Amongst three cysteine point mutants, C36S shows no measurable change in absorbance at 412 nm upon the increasing concentrations of the substrate t-BOOH, i.e., the total free thiol content of this mutant did not change upon increasing the substrate (t-BOOH) concentration. No decrease in the free thiol content in the context of increasing substrate hydroperoxide concentrations reflects the catalytic inactivity of the C36S mutant. This experimental result, along with the result obtained through the multiple sequence alignment, corroborates the fact that without the primary nucleophilic attack by C36, the entire catalytic cycle of SaGpx cannot proceed, explaining the complete loss of its enzymatic activity, which in turn suggests the peroxidative role of C36. In contrast, intriguingly, both C64S and C82S mutants exhibited a decline in total free thiol content of the enzyme upon increasing hydroperoxide concentrations, suggesting that either of them may play the potential role as resolving cysteines. The C64S mutant behaved similarly to the wild type SaGpx. However, C82S mutant displayed a steeper decrease in free thiol content compared to the wild type SaGpx. This DTNB-based assay method relies on tracking free thiol groups to gauge thiol dependent peroxidase function of SaGpx. The curve representing wild type SaGpx got saturated at equimolar concentration (20µM) of substrate tBOOH. The saturated levels of TNB generation could be due to the residual thiol content of the oxidized wild type SaGpx enzyme representing one unreacted thiol group (as three cysteines are there in the wild type, one cysteine will be always available even in the oxidized state of the protein for interaction with DTNB to form TNB) and the fraction of unreacted enzyme molecules that are not taking part in the catalytic conversion even after the presence of hydroperoxide substrate. In case of C64S mutant, the decrease in the saturation point of DTNB titration could be due to the decrease in the residual thiol content, consistent with the mutation of one cysteine to serine. In case of C82S mutant, the increased rate of depletion of the total thiol content could be due to the higher rate of conversion of catalytically active enzyme into the oxidized state involving the formation of disulfide bond between C36 and C64. As a result, the majority of the free thiol groups are getting converted into disulfide species, which may lead to lesser TNB generation upon increasing hydroperoxide concentrations. However, further studies are required to probe the precise roles of these cysteines at 64^th^ and 82^nd^ positions in the catalytic activity of SaGpx. Interestingly, no prominent decreasing pattern was observed in case of Q70A, W124A, N125A mutants as compared to the wild type SaGpx. The loss of the enzymatic activity of these mutants may stem from the disruption of the enzyme’s activity, suggesting their plausible role in substrate and/or active site stabilization rather directly acting as peroxidative or resolving cysteines. Further kinetic experiments were done to clarify the roles of these individual residues (below sections).

### 4. SaGpx utilizes a Thioredoxin-Dependent Redox Cascade

In the canonical glutathione peroxidase (Gpx) pathway, reducing equivalents are provided by glutathione (GSH), which is recycled by glutathione reductase. However, *Staphylococcus aureus* lacks the enzymatic machinery required for GSH biosynthesis, and no homolog of glutathione reductase has been identified in its genome. This suggests that a canonical GSH-dependent peroxidase system is unlikely to operate in this bacterium implying that an alternative redox partner must support SaGpx activity *in vivo*. Previous studies on Gpx homologs, including those from *Populus* [^23^], *Saccharomyces cerevisiae* [^24^], and *Trypanosoma cruzi* [^28^] have shown that these enzymes utilize thioredoxin or other thiol-based reducing agents rather than GSH. These observations raised the possibility that staphylococcal thioredoxins (SaTrx1/SaTrx2), a key redox mediator in *S. aureus*, may act as the alternative electron donor for SaGpx. This hypothesis was validated utilizing the merits of coupled assay. In the coupled assay system, an *in vitro* thioredoxin-dependent redox cascade has been constructed consisting of SaGpx, SaTrx1/2, staphylococcal thioredoxin reductase (SaTR), and NADPH. In this system, electrons should flow sequentially from NADPH to SaTR, from SaTR to SaTrx, and finally from SaTrx to SaGpx through repetitive thiol–disulfide exchange reactions. As substrate hydroperoxide reduction by SaGpx was coupled to NADPH oxidation to NADP^+^, which could be monitored spectrophotometrically at 340 nm, the specific absorbance maximum of NADPH. At the outset, the decline in absorbance at 340 nm upon the complete set up of SaGpx-SaTrx-SaTR electron relay system would thus provide a direct readout of electron transfer through this enzyme cascade and enable evaluation of SaGpx activity under physiologically relevant reducing conditions. For this purpose, we utilized the time-dependent coupled assay. Using this strategy, we first validated that SaGpx accepts electrons from SaTrx1 to reduce tert-butyl hydroperoxide (t-BOOH). The steady decrease **(Fig. S2A)** in NADPH absorbance confirmed the functional coupling of SaGpx to the thioredoxin system, establishing SaTrx1 as its plausible cognate electron donor. However, in contrast, SaTrx2 **(Fig. S2B)** didn’t show a significant decrease in NADPH absorbance as compared to SaTrx1. Further kinetic characterisations were performed to validate the claim of thioredoxin acting as a plausible cognate electron donor of SaGpx. This result not only distinguishes SaGpx from canonical GSH-dependent Gpxs but also places it within the expanding group of thioredoxin-linked peroxidases described in prokaryotes and lower eukaryotes.

### 5. Kinetic characterization of SaGpx mutants shows decrease in catalytic efficiency

Estimation of enzyme kinetic parameters provide critical information about the enzyme activity and catalytic mechanism. The enzyme kinetic parameters of wild type SaGpx and the SaGpx point mutants were calculated using the coupled assay as described in the materials and method section. Initially, this coupled assay was used to screen optimum substrate for wild type SaGpx using different hydroperoxide substrates like hydrogen peroxide (H_2_O_2_), tertiary-butyl hydroperoxide (t-BOOH) and cumene hydroperoxide. The change in absorbance at 340 nm, corresponding to terminal electron donor NADPH consumption per unit time, was recorded. Using the molar extinction coefficient of NADPH (6220 M^−1^·cm^−1^), the Michaelis constant (K_m(app)_) and maximal reaction velocity (V_max(app)_) were calculated by fitting the experimental data to the Michaelis-Menten equation. Turnover number (K_cat(app)_) and catalytic efficiency (K_cat(app)_/K_m(app)_) were subsequently derived to evaluate the substrate specificity of the wild type SaGpx as well as to compare the catalytic efficiency of the wild type and mutant SaGpx. Among the three peroxides tested, SaGpx exhibited the highest affinity for cumene hydroperoxide, reflected by the lowest K_m(app)_ value of 4.78 µM and an apparent catalytic efficiency of 2.1×10^4^ M^−1^·s^−1^ **(Table 1, Fig. 3A)**. Accordingly, cumene hydroperoxide was selected as the substrate for all subsequent enzyme kinetic experiments. *Staphylococcus aureus* has two earlier reported thioredoxin homologs in its genome, SaTrx1 (Uniprot Id: Q2FZD2) and SaTrx2 (Uniprot Id: Q2G000) respectively. Following the screening of the most favoured hydroperoxide substrate, the favoured alternative electron donor of SaGpx was examined using the same coupled assay as described in the materials and method section 3.3. SaGpx exhibited interaction with SaTrx1, yielding a K_m(app)_ value of 2.07 µM and an apparent catalytic efficiency of 1.18×10^5^ M^−1^·s^−1^ **(Table 2, Fig. 3B)**. However, no consistent kinetic behaviour was observed with SaTrx2, suggesting a lack of electron transfer between SaTrx2 and SaGpx.

**Table 1.**
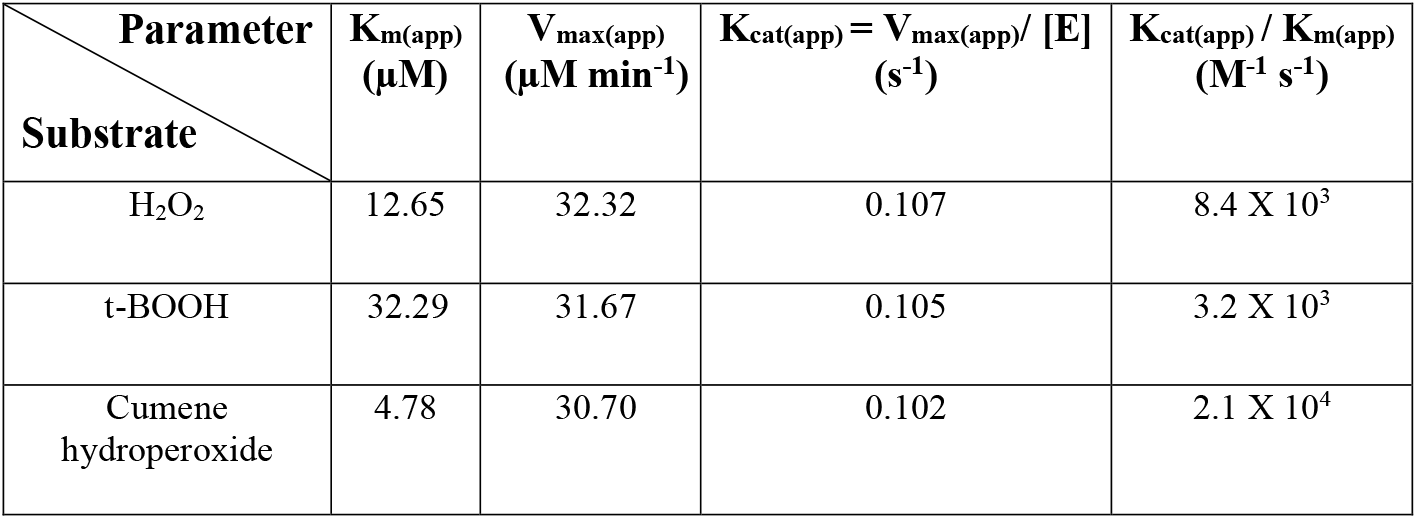
Kinetic parameter estimation of SaGpx with different peroxide substrates. Cumene hydroperoxide exhibits lowest K_m(app)_ of 4.78 µM with highest apparent catalytic efficiency of 2.1×10^4^ M^−1^·s^−1^ compared to H_2_O_2_ and t-BOOH suggesting highest affinity of SaGpx towards cumene hydroperoxide amongst the tested hydroperoxide substrates.

**Table 2.** Kinetic parameter estimation of SaGpx with SaTrx1 and SaTrx2. SaTrx1 exhibits K_m(app)_ of 2.07 µM with apparent catalytic efficiency of 1.18×10^5^ M^−1^·s^−1^ with cumene hydroperoxide as substrate. This confirms SaTrx1 as the alternative electron donor of SaGpx. In contrast, SaTrx2 does not show any electron relay in the coupled assay suggesting no interaction with SaGpx. (ND: Not Detected)

| Parameter<br>Protein | $K_{m(app)}$<br>($\mu\text{M}$) | $V_{max(app)}$<br>($\mu\text{M min}^{-1}$) | $K_{cat(app)} = V_{max(app)} / [E]$<br>($\text{s}^{-1}$) | $K_{cat(app)} / K_{m(app)}$<br>($\text{M}^{-1} \text{s}^{-1}$) |
| --- | --- | --- | --- | --- |
| SaTrx1 | 2.07 | 73.53 | 0.245 | $1.18 \times 10^5$ |
| SaTrx2 | ND | ND | ND | ND |

**Figure 3.**
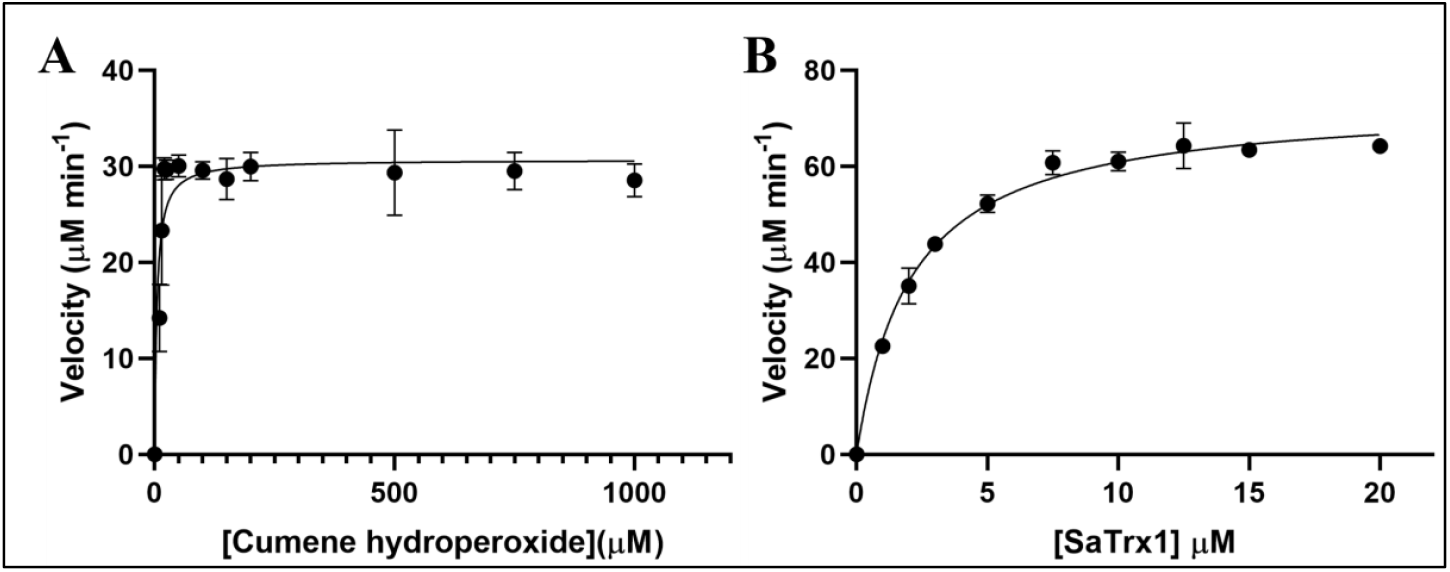
Michaelis-Menten kinetic characterization of SaGpx with Cumene hydroperoxide as a substrate and SaTrx1 as an electron donor. (3A) Enzyme kinetic analysis of SaGpx with the favoured substrate cumene hydroperoxide. (3B) Enzyme kinetic analysis of SaGpx with the plausible cognate electron donor SaTrx1. Reaction velocity was plotted against substrate concentration (3A) and cofactor (SaTrx1) concentration (3B) and fitted into Michaelis-Menten equation [V=V_max_*S/(k_m_+S)]. Different kinetic values are listed in Table 1 and 2.

To gain deeper insight into the precise role of the highly conserved amino acid residues in SaGpx mediated enzyme catalysis, the coupled assay was performed for all the selected SaGpx point mutants using SaTrx1 as the cognate electron donor and cumene hydroperoxide as the substrate. Kinetic parameters were subsequently determined for the selected SaGpx mutants to elucidate the specific contributions of these residues to enzyme function. The mutant C36S displayed no detectable electron relay from NADPH (behaves like SaGpx negative control experiment), indicating complete functional disruption of this mutant. The loss of enzymatic function of C36S is also supported by the result obtained from the DTNB-based hydroperoxide titration assay **(Fig. 2B)** and suggests C36 may play the role of peroxidative cysteine. Intriguingly, amongst C64S and C82S point mutants, the C64S variant retained comparable catalytic activity to the wild type SaGpx, with a K_m(app)_ of 5.33 µM and an apparent catalytic efficiency of 1.8×10^4^ M^−1^·s^−1^ **(Table 3, Fig. S3)**, indicating C82 could play the role of resolving cysteine. In the contrary, unlike the results found in DTNB based hydroperoxide titration assay, the C82S mutant shows no electron relay from NADPH in the coupled assay. This could be due to the unresponsiveness of reduced SaTrx1 towards the oxidized SaGpx having disulfide bond between C36 and C64. Perhaps, it may suggest that the C36-C64 disulfide-bonded oxidized SaGpx may need an alternative electron donor other than SaTrx1 to resolve its intramolecular disulfide bond. Studies on the glutathione peroxidase, Hyr1 from *Saccharomyces cerevisiae* suggests the involvement of different reducing agents to recycle the oxidized glutathione peroxidase in varying oxidative stress conditions [^24^]. This reference also suggests the possibility of involvement of a different reducing agent to resolve the disulfide bond formed between C36 and C64. However, further studies in this direction are required to elucidate the precise role of the C64 in the enzymatic activity of SaGpx. Interestingly, the non-cysteine substitutions significantly compromised the catalytic efficiency of SaGpx. The Q70A, W124A, and N125A mutants displayed markedly elevated K_m(app)_ values for the substrate cumene hydroperoxide (364.3 µM, 359.4 µM, and 10340 µM, respectively) coupled with drastic reductions in apparent catalytic efficiency (0.04×10^4^, 0.04×10^4^, and 0.003×10^4^ M^−1^·s^−1^). These kinetic impairments emphasize their importance in ensuring proper substrate binding and turnover rather than directly mediating the thiol oxidation/reduction. Notably, the severe defect observed in the N125A mutant underscores its indispensable role in catalysis, consistent with reports of similar disruptions in enzyme function upon asparagine-to-alanine substitution in *Drosophila melanogaster* glutathione peroxidase [^29^]. Collectively, these findings confirm C36 as the peroxidative cysteine, and C82 as the resolving cysteine, and establish key non-cysteine residues as crucial for substrate stabilization and efficient peroxide detoxification in SaGpx. However, to probe the exact role of C64 in catalysis, further studies are needed.

**Table 3.** Comparison of Kinetic parameters between different mutants of SaGpx. C36S shows no detectable change in NADPH absorbance suggesting functional disruption of protein due to this mutation. C64S shows similar kinetic parameters as of the wild type SaGpx, suggesting the role of C82 as resolving cysteine. However, C82S does not exhibit any electron relay from NADPH. The possible reason for this could be the unresponsiveness of the SaTrx1 towards the disulfide bond formed between C36 and C64. The non-cysteine mutants, Q70A, W124A, and N125A shows significantly elevated K_m(app)_ values (364.3 µM, 359.4 µM, and 10340 µM, respectively) coupled with drastic reductions in apparent catalytic efficiency (4.4×10^2^, 4.2×10^2^, and 3.0×10^1^ M^−1^·S^−1^), suggesting their role in substrate binding and turnover rather than directly mediating thiol oxidation. (ND: Not Detected)

| Parameter<br>Variants | $K_{m(app)}$<br>(μM) | $V_{max(app)}$<br>(μM $min^{-1}$) | $K_{cat(app)} = V_{max(app)} / [E]$<br>($s^{-1}$) | $K_{cat(app)} / K_{m(app)}$<br>($M^{-1} s^{-1}$) |
| --- | --- | --- | --- | --- |
| Wild type | 4.78 | 30.70 | 0.102 | $2.1 \times 10^4$ |
| C36S | ND | ND | ND | ND |
| C64S | 5.33 | 29.74 | 0.099 | $1.8 \times 10^4$ |
| C82S | ND | ND | ND | ND |
| Q70A | 364.3 | 48.87 | 0.162 | $4.4 \times 10^2$ |
| W124A | 359.4 | 45.66 | 0.152 | $4.2 \times 10^2$ |
| N125A | 10340 | 94.34 | 0.314 | $3.0 \times 10^1$ |

Q70A, W124A and N125A mutants shows ∼76-fold, ∼75-fold and ∼2100-fold increase in the substrate K_m(app)_ value than the wild type SaGpx. The increased K_m(app)_ values of Q70A, W124A and N125A mutants suggests their role in the substrate binding. Another kinetic parameter, K_cat(app)_ represents the turnover number of an enzyme, which implies the maximum number of substrate molecules converted by a single enzyme into products per unit time. Importantly, the experimentally obtained K_cat(app)_ values of the wild type SaGpx did not change drastically in case of the tested point mutants. The comparable K_cat(app)_ values of the tested point mutants to the wild type SaGpx suggest that, once the substrate is bound, the rate of conversion from substrate to product is similar for the wild type SaGpx and all the tested point mutations. Therefore, the tested point mutations might not impact the formation of the substrate transition state to the extent that they do adversely impact the binding of the incoming hydroperoxide substrate. However, the catalytic efficiency calculation, represented by K_cat(app)_/K_m(app)_ provides a more composite picture of an enzymes fitness in comparison to considering K_m(app)_ and K_cat(app)_ values alone. Comparing catalytic efficiency of wild type SaGpx enzyme with its different tested mutant versions provides a holistic comparison of their individual role.

### 6. Spectroscopic analysis of SaGpx structural features

#### 6.1. CD spectroscopy-based analysis reveals structural transition of SaGpx upon oxidation

Superimposition of the reduced (C36 mutant) and the modelled oxidized state of SaGpx shows the complete unwinding of the alpha helix harbouring the putative resolving cysteine at position 82 in the oxidized model. To experimentally validate this structural shift in transition from reduced to oxidized state, circular dichroism (CD) spectroscopy was employed under native and oxidizing conditions using 50 µM tert-butyl hydroperoxide (t-BOOH). The CD spectra show deconvolution of the characteristic peak of alpha Helix at ∼222nm **(Fig. S4A)** indicating loss of helical content. This spectral deconvolution supports the predicted structural rearrangement, confirming the oxidation induces unwinding of the α-helix, consistent with the homology-based structural model of oxidized SaGpx **(Fig. S5)**.

#### 6.2. Fluorescence-based assessment of active site microenvironment in SaGpx

Sequence and structural analysis of the SaGpx indicates a potential catalytic role for tryptophan, located within the enzyme’s active site. Given the sensitivity of tryptophan’s indole fluorescence towards its surrounding environment, intrinsic fluorescence spectroscopy was employed to monitor conformational changes during redox transition. Upon exposure to tert-butyl hydroperoxide (t-BOOH), a hyperchromic shift in the emission spectra was observed **(Fig. S4B)**, suggesting alterations in the local environment of the tryptophan residue. In the reduced state, tryptophan is closely positioned near polar residues such as cysteine and glutamine, whose sulfhydryl and amide groups are known to quench tryptophan fluorescence [^33^]. However, in the oxidized form, structural rearrangement, specifically the unwinding of the α-helix, results in displacement of these quenching residues away from tryptophan **(Fig. S5C)**. This spatial reorganization reduces quenching and leads to enhanced fluorescence intensity, consistent with the observed spectral shift and supporting the modelled conformational transition from reduced to oxidized states.

#### 6.3. Spectroscopic validation of sulfenic acid intermediates of SaGpx under oxidative condition

The detection of sulfenic acid, an oxidized form of cysteine, is crucial in understanding the catalytic mechanism of SaGpx, a peroxidase enzyme [^34^]. The catalytic cycle of SaGpx starts with the nucleophilic attack of the thiolate anion (S^−^) of the peroxidative cysteine to the peroxyl bond of the substrate peroxide. This initial attack results in the formation of sulfenic acid (S-OH), a key intermediate in the peroxidase cycle [^35^]. Sulfenic acid formation was traced by employing a spectroscopic approach utilizing NBD-Cl (4-Chloro-7-nitrobenzofurazan). NBD-Cl is capable of binding to both thiol groups and sulfenic acid moieties but forms distinct adducts with unique spectral properties for each. The thiol adducts exhibit a characteristic absorbance peak at 420 nm, whereas the sulfenic acid adducts show a distinct peak at 347 nm [^36^]. C64S+C82S double mutant of SaGpx was used in this sulfenic acid detection assay to ensure the observed sulfenic acid is coming only from peroxidative cysteine and not influenced by other cysteines. In the presence of substrate hydroperoxide (t-BOOH), the peroxidative cysteine gets converted into sulfenic acid. As there was no cysteine in the protein itself and no external reducing agents were added in the reaction mixture, the sulfenic acid could not be resolved into either disulfide bond or cysteine thiol. The sulfenic acid thus formed forms sulfenic acid adduct when incubated with NBD-Cl. The presence of the characteristic peak (at 347 nm) for sulfenic acid was observed in the oxidized protein sample while no peak at 420 nm **(Fig. S4C)**. However, no such peak observed in reduced protein sample at 347 nm. Reduced protein sample shows the characteristic peak (at 420 nm) for thiol adduct of NBD-Cl. Comparison of the peaks confirms the formation of sulfenic acid intermediate during interaction with substrate hydroperoxide.

### 7. Crystal structure of C36S SaGpx mutant

#### 7.1. Overall quality of the crystal structure

The crystal structure of C36S SaGpx mutant has been solved at 1.5 Å. The structure was solved in C-centred orthorhombic space group C 2 2 2_1_ with unit cell parameters of, a = 62.79 Å, b = 95.92 Å, c = 59.78 Å and α =β =γ = 90º. The crystal structure of C36S SaGpx mutant was solved as monomer. The asymmetric unit contains one SaGpx protomer with Matthew’s coefficient of 2.48 Å^3^/Da corresponding to the calculated solvent content of 50.53% in the unit cell. No significant oligomeric structural assembly could be formed using the symmetry mates of the C36S SaGpx monomer in the asymmetric unit. The structure has been solved by molecular replacement using the atomic coordinates of previously solved x-ray crystal structure of the poplar glutathione peroxidase 5 in the reduced form (PDB Id: 2P5Q) [^23^] which shows overall 46.02% amino acid sequence identity with SaGpx amino acid sequence upon NCBI BLAST. In the crystal structure, the asymmetric unit shows a monomeric form of SaGpx with a subunit molecular mass of 18.2 kDa, with clear electron density for all 158 amino acid residues. The diffraction data exhibited completeness of 99.86% and multiplicity of 2.0, with an overall I/σ(I) ratio of 3.38 in the highest resolution shell. The final refined model structure of C36S mutant shows R_work_ and R_free_ factors of 0.1505% and 0.1701% respectively, reflecting excellent agreement between the constructed model and the experimental data. The final refined structure exhibits 98.72% of amino acid residues in the most favoured regions of the Ramachandran plot with no residues in disallowed regions and no rotamer outliers, confirming the stereochemical validity of the model. The detailed information of the crystal diffraction data collection, scaling and refinement is summarised in Table S1.

#### 7.2. Overall Structure

Overall, the structure of C36S SaGpx mutant was found to be globular with dimensions of ∼ 31.2 Å X 36.8 Å X 38.2 Å in length, width and height. The crystal structure of C36S SaGpx mutant displays the well documented glutathione peroxidase fold [^37^]. The reduced structure **(Fig. 4A-C)** is comprised of seven β-strands and four α-helices along with some additional secondary structural elements. The secondary structure starts with two 3_10_ helices (α1 and α2) which are separated by two β-strand (β1 and β2) forming a β-hairpin. Other β-strand (β3 to β7) forms the central twisted β-sheet by arranging themselves in parallel (β3-β4-β5) and antiparallel (β3-β6-β7) sections. The centrally twisted β-sheet is surrounded by three α-helices (α3, α4 and α7) on one side and one α-helix (α5) on the other side. Helices α3 and α7 are parallel to each other and their axes are nearly parallel to the central β-sheet. Helices α4 and α5 are perpendicularly oriented to each other and to the helices α3 and α7. The loop between α5 and β6 harbours a 3_10_ helix (α6). The average B factor of 27.55 indicates that the monomeric globular structure of C36S SaGpx to be stable. However, the relative distribution of B factors demonstrates higher structural vibrations of the loop connecting α6 and β6. Intriguingly, this region has been found to be extended in human Gpx1 and was shown to be important for its stringent substrate specificity and tetramerization. In SaGpx this region is structurally reminiscent to human Gpx4 in terms of its length and structural flexibility and thereby may also confer the broader substrate specificity of SaGpx. The structure of C36S SaGpx mutant shows clear side chain electron densities of the sulfhydryl (-SH) groups of the amino acid positions at C64 and C82 **(Fig. S6)**. The catalytic attenuation of the C36S mutant **(Fig. 2B and Table 3)** as well as the absence of any further disulfide bond(s) in the C36S SaGpx mutant structure ensures the reduced conformational state of the protein. However, wild type SaGpx could not be successfully crystallized in its oxidized state. Therefore, the oxidized model structure **(Fig. S5A, S7)** of wild type SaGpx was obtained from homology modelling using Modeller software [^38^]. The crystal structure of the poplar glutathione peroxidase 5 in the oxidized form (PDB Id: 2P5R) [^23^] has been used as the template for homology modelling. The oxidized model shows a disulfide bond formed between cysteines at 36^th^ and 82^nd^ position. The formation of the disulfide bond in between C36 and C82 is also supported by the enzyme kinetics data that shows, C64S SaGpx mutant catalytically behave similarly as wild type SaGpx, whereas C36S SaGpx and C82S SaGpx point mutants were found to be catalytically inert. Superimposition **(Fig. S5B)** of the reduced (obtained from crystal structure) and the oxidized (obtained from homology modelling) forms of SaGpx shows a Root-Mean-Square-Deviation (RMSD) of 1.095 Å (based on the alignment of 121 C_α_ positions). This high RMSD value is largely due to the unwinding of a complete α-helix (helix α4) into a loop during the transition from reduced to oxidized state.

**Figure 4.**
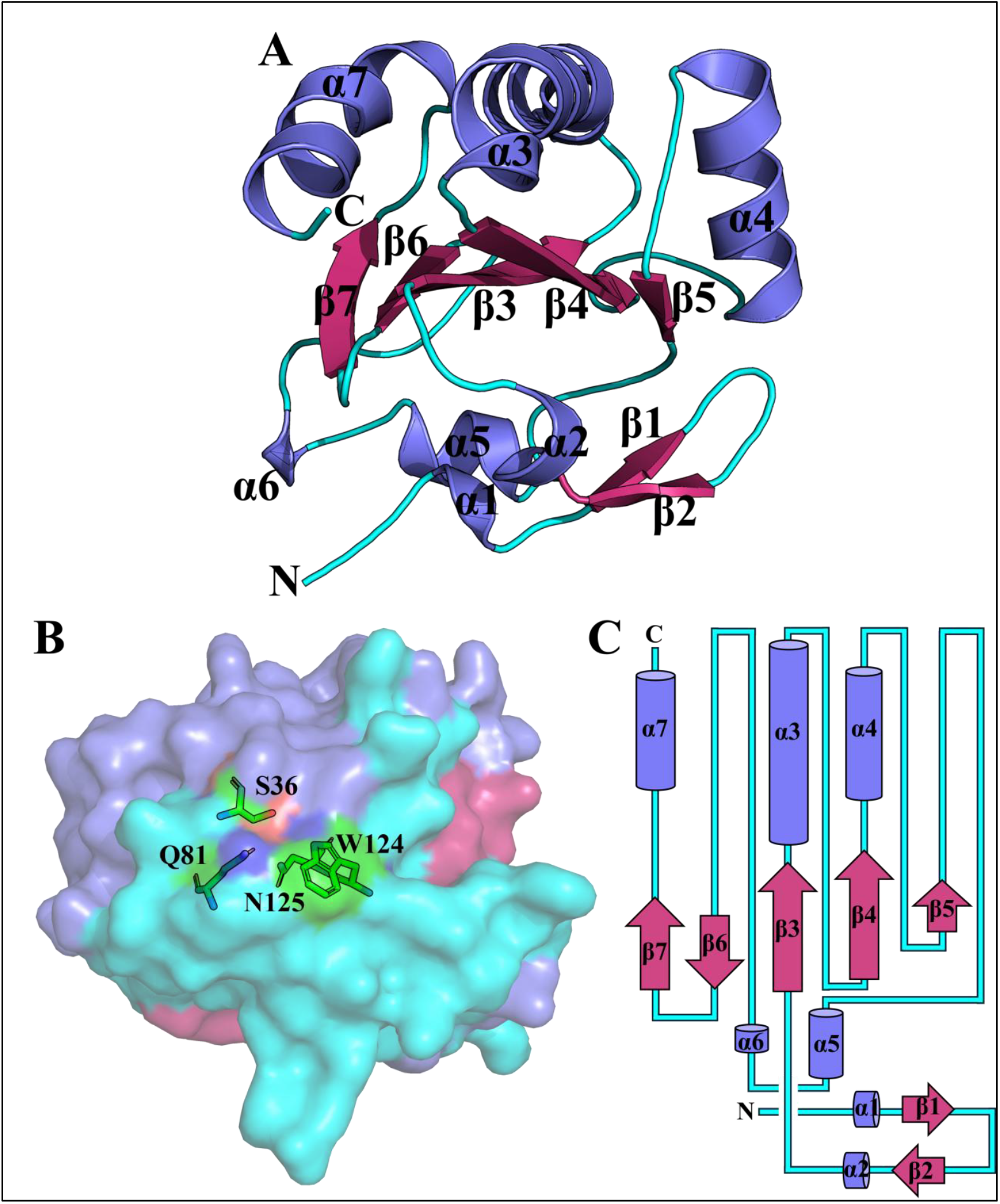
Overall structural feature of C36S SaGpx mutant. (4A) The crystal structure of C36S is represented as cartoon. The structure shows Gpx-fold consisting of α-β-α sandwich, where centrally aligned β-strands (red) are flanked by α-helices (blue) with additional short secondary structural elements at the N-terminal end. (4B) Overall surface representation highlighting the active site residues, forming the catalytic tetrad. (4C) Topology diagram of the crystal structure of C36S SaGpx showing relative structural disposition of the secondary structural elements. Coloured structural elements represent the presence of the Gpx fold: α helical contents were represented as blue, β strands were represented as red and loops as cyan lines.

### 8. Structural comparison of SaGpx with Gpx homologs

Gpx homologs with more than overall 40% amino acid sequence identity with SaGpx amino acid sequence were identified by BLASTp search against PDB database; moreover, previously solved crystal structures with RMSD values below 1.5 Å from DALI structural alignment [^39^] with C36S SaGpx crystal structure were selected for comparison with SaGpx. Multiple sequence alignment of the amino acid sequences **(Fig. 1)** revealed that SaGpx lacks an extended N-terminal amino acid sequence observed in certain mammalian Gpx counterparts such as human Gpx3, human Gpx5, and human Gpx6, and human Gpx7. This extended N-terminal region in human isoforms is generally associated with signal peptide function. Whereas human Gpx8 contains an N-terminal transmembrane domain [^40^]. Additionally, SaGpx lacks an internal stretch of around 15 amino acids, labelled here as loop1 **(Fig. 1)**, which is present in tetrameric human Gpx homologs (human Gpx 1,2,3,5,6). Structural superimposition with one such representative human Gpx isoform, Gpx1 (PDB Id: 2F8A) **(Fig. 5A)**, containing loop 1, confirmed its absence in SaGpx crystal structure. Interestingly, this surface-exposed loop appears to lie in front of the active site and partially occludes W124, one of the highly conserved amino acids of the Gpx catalytic tetrad, present in the active site. The position of this loop is likely the reason for restricting access to the complex lipid hydroperoxides, thereby limiting the substrate specificity of some human Gpx isoforms (Gpx isoforms 1, 2, 3, 5, 6) [^31^]. In contrast, the extensively studied human Gpx4 **(Fig 7B)**, which also lacks loop 1, just like SaGpx, exhibits broader substrate specificity and efficiently reduce complex lipid hydroperoxides [^31^]. Given that SaGpx shares more than 40% amino acid sequence identity (overall) with human Gpx4 and its crystal structure is also structurally reminiscent of human Gpx4 (RMSD of 0.62 Å with 126 Cα positions), it is plausible that SaGpx may also process complex lipid peroxides, although further experimental confirmation is required. A further distinction is the absence of a stretch of 5 amino acids labelled as loop2 (**Fig. 5A)** at the end of the helix 4 in SaGpx. In human Gpx1, Gpx2, Gpx3, Gpx5, and Gpx6, this additional segment displaces helix α4 towards the protein surface as evident from the structural superimposition. This loop1 and loop2 present in the interface of tetrameric human Gpx isoforms and are responsible for their higher order oligomerization [^31^]. Also, the superimposition of SaGpx with human Gpx7 (PDB ID: 2P31), which is present in monomeric form shows overall structural similarity **(Fig. 5B)** with an RMSD value of 0.77 Å (based on the alignment of 137 Cα positions). The absence of extra amino acid residue at the end of helix 4 (loop2) and loop1 may be considered as the structural justification for the monomeric form of SaGpx.

**Figure 5.**
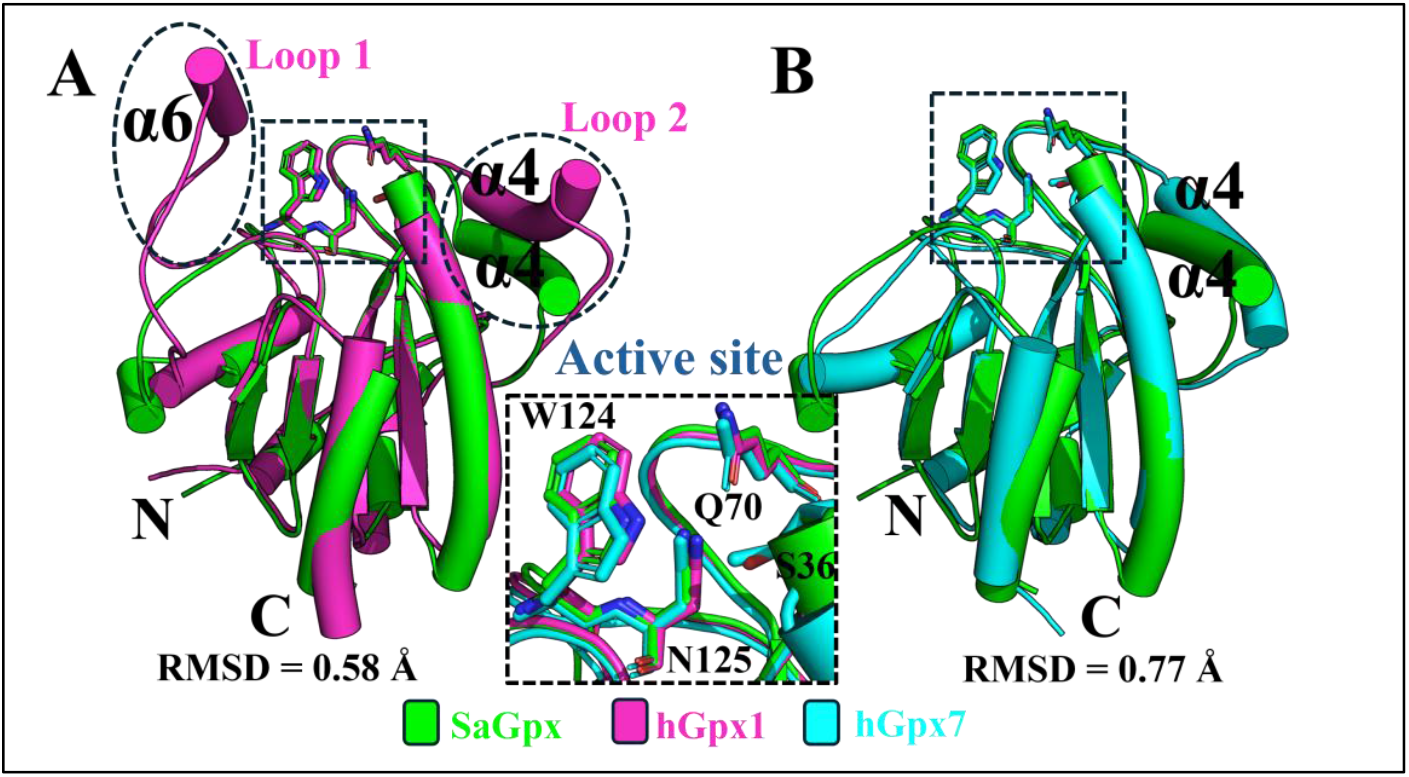
Structural comparison of SaGpx with other glutathione peroxidase homologs. (5A) Structural superimposition of SaGpx (green) with tetrameric human Gpx1 (PDB ID: 2F8A, magenta). Black dotted circles highlight oligomerization loops (loop1 and loop2) present in human Gpx1 but absent in SaGpx. (5B) Structural superimposition of SaGpx (green) with monomeric human Gpx7 (PDB ID: 2P31, cyan). The superimposition exhibits similar absence of oligomerization structural elements in both proteins, confirming the monomeric nature of SaGpx. Inset: The central panel (black dotted box) shows an enlarged view of the conserved active site architecture across both homologs.

### 9. Active site architecture

The high-resolution electron density map (2Fo-Fc map contoured at 1σ) revealed well-defined active site residues of C36S SaGpx mutant **(Fig. 6A-B)**. Structural alignment with known glutathione peroxidase crystal structures (with RMSD values less than equal to 1.5 Å) confirms that the active site residues, C36, Q70, W124, and N125 are spatially well conserved in SaGpx and together constitute the catalytic tetrad of the enzyme. The peroxidative cysteine (C_P_) residue at the 36^th^ position (as S36 in the described crystal structure of the recombinant protein corresponds to the C36 of the native protein, further discussion is continued with considering C36 to avoid any confusion with the native condition) is the first cysteine residue starting from the N-terminal end of the protein. The 3D structural location of C36 is conserved throughout the Gpx homologs (although cysteine is replaced by selenocysteine in case of eukaryotic Gpx homologs) and lies in the loop connecting the third β-strand to third α-helix. Sequence analysis further demonstrated that this residue falls within the canonical Gpx signature motif 1 (G[K/R]x[L/V][I/L]I[V/E/T]NVA[S/T/A][E/Q/L/Y][**C/U**]G[L/T]T) **(Fig. 1)**. The neighbouring residues like G37, F38, and T39 stabilizes the C36 backbone by forming hydrogen bonds. The crystal structure of SaGpx shows the possible resolving cysteines, C64 and C82 are located away (∼ 11 Å and ∼13 Å respectively) from the peroxidative cysteine, C36. The 64^th^ cysteine is part of the conserved Gpx signature motif 2. The 82^nd^ cysteine is present in a non-aligned position as evident from multiple amino acid sequence alignment studies, however, upon structural comparison with thioredoxin-dependent 2-cysteine Gpx isomers like poplar Gpx and yeast Gpx, this residue is spatially conserved. The superimposition of SaGpx with reported reduced structures of human Gpx4 (PDB Id: 2OBI, RMSD of 0.62 Å based on alignment of 126 Cα atoms), poplar Gpx (PDB Id: 2P5Q, RMSD of 0.71 Å based on alignment of 133 Cα atoms) and *S. mansoni* Gpx (PDB Id: 2V1M, RMSD of 0.62 Å based on alignment of 134 Cα atoms) suggest that the crystal structure of C36S SaGpx mutant exists in its reduced form. The remaining active site residues of the catalytic tetrad, Q70, W124 and N125 are positioned near the thiol group (-SH) of the peroxidative cysteine (in the C36S mutant, near the hydroxyl group of S36) consistent with their role in lowering the pKa of the peroxidative cysteine and thereby helps to maintain in its reactive thiolate anion (-S^−^) form at physiological pH (7.4). These residues also play the crucial role in binding of the peroxide substrate and preparing it for subsequent enzymatic reaction as evident from biochemical characterization **(Table 3)**. This nucleophilic state of sulfur is necessary for the nucleophilic attack on the electron deprived oxygen of the incoming peroxide substrate. In the crystal structure, a water molecule is present within the hydrogen bonding distance of the key interacting atoms of the catalytic tetrad residues (Oξ1 of Q70, Nξ1 of W124 and Nδ2 of N125). Interestingly, a symmetry-related leucine residue (L158) was observed with its carboxyl oxygen in the carboxylate terminal (the OXT atom) directed towards this water molecule. The spatial arrangement of the OXT atom and the nearby water molecule, together, may resemble the geometry of a peroxide functional group, suggesting a structural mimic that may reflect how the active site accommodates peroxide substrates **(Fig. 6A-B)**. The catalytic residues exhibited uniformly low B-factors (19–24 Å^2^), indicating high local stability of the SaGpx active site. Importantly, the comparable B-factors of the water molecule and the OXT atom to those of the catalytic tetrad residues further support the notion that the peroxide substrate would remain structurally stabilized during the interaction with the active site residues. This special structural arrangement, elicited by two symmetry-related C36S SaGpx monomers, is highly stable and was found to be refractile during soaking the crystal with different hydroperoxide substrates at low to mid millimolar concentrations, with no traceable electron densities for the incoming hydroperoxides. However, soaking the same crystal with very high hydroperoxide concentrations resulted in crystal melting with no visible diffraction spots, which may occur due to the disruption of the crystal contacts overpowered by competitive hydroperoxide binding.

**Figure 6.**
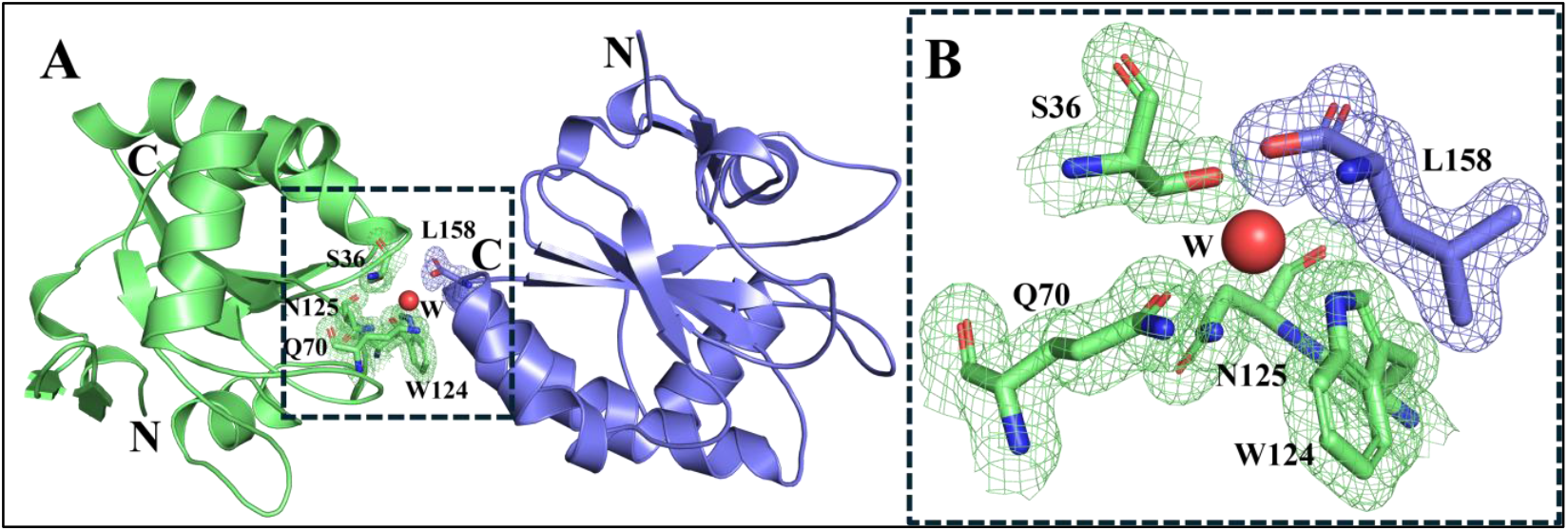
Active site architecture and the catalytic environment of SaGpx. (6A) Spatial organization of the active site amino acid residues. The three-dimensional arrangement shows a symmetry-related SaGpx monomer (colored blue) with its C-terminal carboxylate (OXT of L158) positions itself within the active site of the neighbouring monomer (colored green) in a different asymmetric unit, hence mimicking the geometry of a hydroperoxide substrate and provides the structural glimpse of potential substrate binding mode of SaGpx. (6B) Magnified view of catalytic residues with electron density validation. High-resolution representation of key catalytic residues supported by electron density maps (2Fo-Fc map contoured at 1σ), demonstrating the well conserved active site architecture.

The active site region of SaGpx is devoid of any definite substrate binding pocket. The catalytic residues are exposed to the surface and cover an area of around 147.51 Å^2^. However, the peroxidative cysteine is present in a positively charged dent, majorly contributed by catalytic tetrad residues **(Fig. 7E)**. Just beneath the active site, there is a large patch of positively charged region. This cationic patch forms a cradle spanning an area of ∼2045 Å^2^. K123, K139, R140, K145, K146, R152 are the major contributors of the positive charge to this patch **(Fig. 7F)**. Two flexible side chains, F118 and Q144, flank this region and display elevated B-factors (55–75 Å^2^ and 21–51 Å^2^, respectively). Their dynamic behaviour suggests a potential gating role that may facilitate the binding of bulky anionic substrates such as phospholipid hydroperoxides. Similar cationic surface patches have been reported in human phospholipid Gpx (Gpx4) [^41^] **(Fig. 7A)** and in *Schistosoma* Gpx [^29^], where they mediate interactions with complex phospholipid substrates. This cationic patch with these two dynamic side chains seems to act as a ‘flycatcher’. By this structural analogy, the SaGpx may function as a bacterial phospholipid glutathione peroxidase. However, although poplar Gpx exhibits more than 40% sequence similarity and a RMSD of 0.71 Å (based on 133 Cα atoms), lacks this cationic patch. Poplar Gpx exhibits a highly negative charged surface regions **(Fig. 7C)** compared to SaGpx. Compared to SaGpx and human Gpx4, the high abundance of negatively charged surface of the Poplar Gpx may confer its role as a heavy metal ion (Cd^2+^) sink, suggesting enzymes’ adaptations to substrate specificity [^23^].

**Figure 7.**
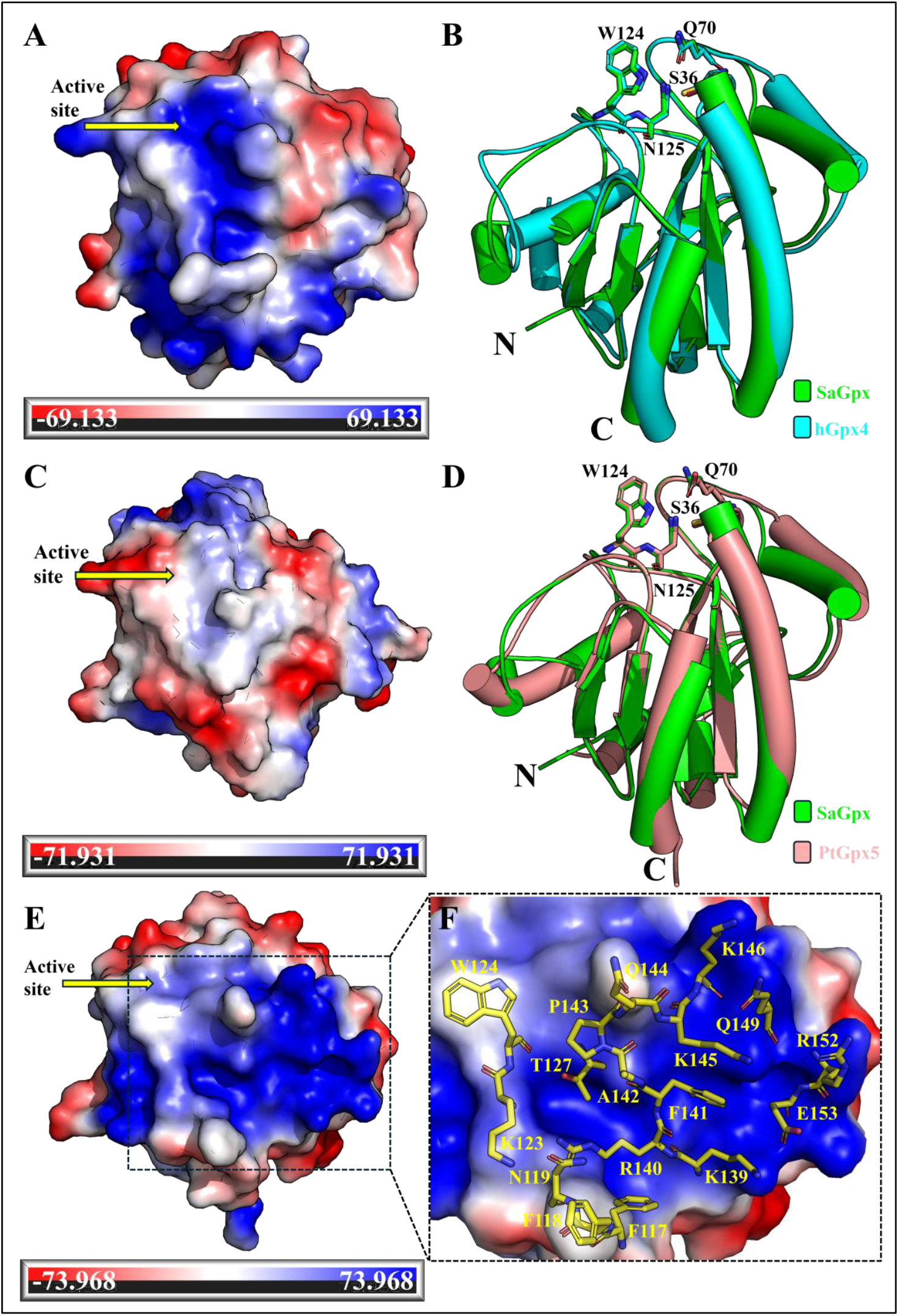
Active site proximal electrostatic surface potential comparison of SaGpx with other Gpx homologs. Comparative analysis shows active site proximal surface electrostatic potentials of (7A) human Gpx4, (7C) poplar Gpx5, and (7E,7F) SaGpx with detailed cationic patch visualization. Structural overlays demonstrate conservation between SaGpx and (7B) human Gpx (cyan) and (7D) poplar Gpx5 (rose). SaGpx resembles human Gpx4 by showing a characteristic cationic patch beneath the active site, whereas poplar Gpx5 presents predominantly negatively charged surface. These electrostatic differences likely reflect the functional optimization for distinct hydroperoxide substrate.

## Discussion

The structural and biochemical analyses, presented herein, reveal that SaGpx represents a distinct class of glutathione peroxidase different from the classical glutathione peroxidase. Though SaGpx exhibits clear structural and functional similarities to canonical mammalian glutathione peroxidases, it differs fundamentally in its catalytic mechanism and electron donor specificity. Classical mammalian Gpx enzymes utilizes selenocysteines as the peroxidative residue for substrate hydroperoxide reduction [^12^]. In contrast, SaGpx uses two cysteine residues for catalysis: a peroxidative cysteine (C_P_) and a resolving cysteine (C_R_). The choice between cysteine and selenocysteine in the active site has a significant effect on how these enzymes interact with their reducing partner during catalytic regeneration. Enzymes with selenocysteine, like human Gpx 4, evolved to use two molecules of glutathione for their reduction and reactivation [^42^]. On the other hand, the two-cysteine variants of Gpx homologs like SaGpx which are found in lower eukaryotes, plants and bacteria prefer thioredoxin or tryparedoxin (proteins characterized by the -CXXC-motif) proteins as reductants [^24,28^]. Based on our enzyme activity assays and site-directed mutagenesis of the catalytic cysteines, SaGpx appears to follow catalytic mechanism similar to atypical 2-cysteine peroxiredoxins. [^43^]

Both Gpx and atypical 2-Cys Prx share the same overall parent fold, that is the thioredoxin (Trx) fold [^37^], composed of centrally aligned 4 β strands flanked by 3 α helices. However, SaGpx like other homologs adopt more precisely the glutathione peroxidase (Gpx) fold which is distinct from the Trx fold in terms of two extra β-strands at the N-terminus along with insertion of one β-strand and one α-helix. In the redox centre of classical Gpx, the peroxidative SeCys is present in the hydrogen bonding distance from a glutamine, a tryptophan, and an asparagine residue, all together form the conserved catalytic tetrad. [^44^] In contrast, the peroxidative cysteine in atypical 2-Cys Prx is present in the hydrogen bonding distance from a threonine and an arginine, collectively form the catalytic triad. [^42^] SaGpx also displays same overall fold as of classical Gpx with sequentially and spatially conserved active site residues (composed of C36, Q70, W124, and N125) which distinctly distinguishes SaGpx from the atypical 2-Cys Prxs. Also, despite of sharing similar parent fold with atypical 2-Cys Prxs and involvement of a second cysteine acting as resolving one, both Gpx and atypical 2-Cys Prx share very little sequence as well as structural similarities with a RMSD of 3.76 Å (based on the alignment of 17 C_α_ positions of Staphylococcal atypical 2-Cys Prx, PDB Id: 3p7x) **(Fig. S9)**. Thus, based on the structural and biochemical evidence reported herein, SaGpx, although being annotated as glutathione peroxidase may be re-classified and included in the class of Gpx-type thioredoxin peroxidase or thioredoxin-dependent Gpx.

The success of the catalytic mechanism of the SaGpx largely depends on the formation and stability of the thiolate anion (-S^−^) of the peroxidative cysteine, which reduces the peroxide bond (-O-O-) through SN2 reaction by acting as a nucleophile. The significant changes in the kinetic parameters of the mutant variants of the catalytic tetrad residues highlights their indispensable role in enzyme activity and substrate stabilization. Considering their roles in catalysis, a possible scheme of the SaGpx catalysis is proposed here **(Fig. 8)**. In the reduced state, the residues Q70, W124 and N125 are coordinated with the C_P_ thiol through hydrogen bond formation. N125 due to the low dielectric constant of its hydrophobic environment, strongly attracts the proton from the C_P_ thiol. Oξ1 of Q70 on the other hand also attracts the proton from the thiol group. The combined pulling force results in the lowering of the pKa of the C36 thiol and formation of the thiolate anion, which is stabilized by the positive charge conferred by the Nξ1 of W124. This nucleophilic form of thiol (thiolate, −S^−^) is ready to attack the substrate hydroperoxide. But to get attacked by the thiolate, the hydroperoxyl group needs to be polarized and stabilized, which is accomplished by the side chains of both Q70 and N125. It could be likely that Nξ2 of Q70, being exposed to the surface help substrate hydroperoxide to identify the active site through hydrogen bonding and guide the substrate to enter the active site where Oξ1 of Q70 and Nδ2 of N125 interact with terminal −OH group of the substrate hydroperoxide in such a way that the Nξ2 of Q70 ‘trap’ the substrate, and the Oξ1 of Q70 and Nδ2 of N125 ‘fix’ or ‘stabilize’ the substrate hydroperoxide, thereby positioning the substrate near the thiolate anion. N125 also stabilizes Q70 by forming hydrogen bond. The involvement of N125 in thiolate anion generation as well as stabilization of substrate makes this residue highly indispensable for SaGpx catalysis. This hypothesis is corroborated by the steep increase (∼100 times) in the K_m_ of N125A SaGpx mutant variant compared to the wild type SaGpx. The Nξ2 of Q70 donates a proton to the proximal oxygen of the hydroperoxide substrate leading to release of ROH and generation of sulfenic acid (S-OH). The sulfenic acid thus formed is resolved by the resolving cysteine with the formation of intramolecular disulfide bond. Transition from the reduced to the oxidized state, renders a local structural change corresponding to the complete unwinding of an α-helix into loop. This secondary structural change confers oxidized SaGpx noticeable structural protrusion, as obtained from the CD and fluorescence based structural changes of SaGpx upon hydroperoxide substrate addition **(Fig. S5D)**. The conformation of the oxidized form of the protein may help the selective docking with the reducing partner protein like thioredoxin. Intriguingly uncoupling of the oxidation-induced structural change of SaGpx may be crucial in designing precise inhibitors against this enzyme.

**Figure 8.**
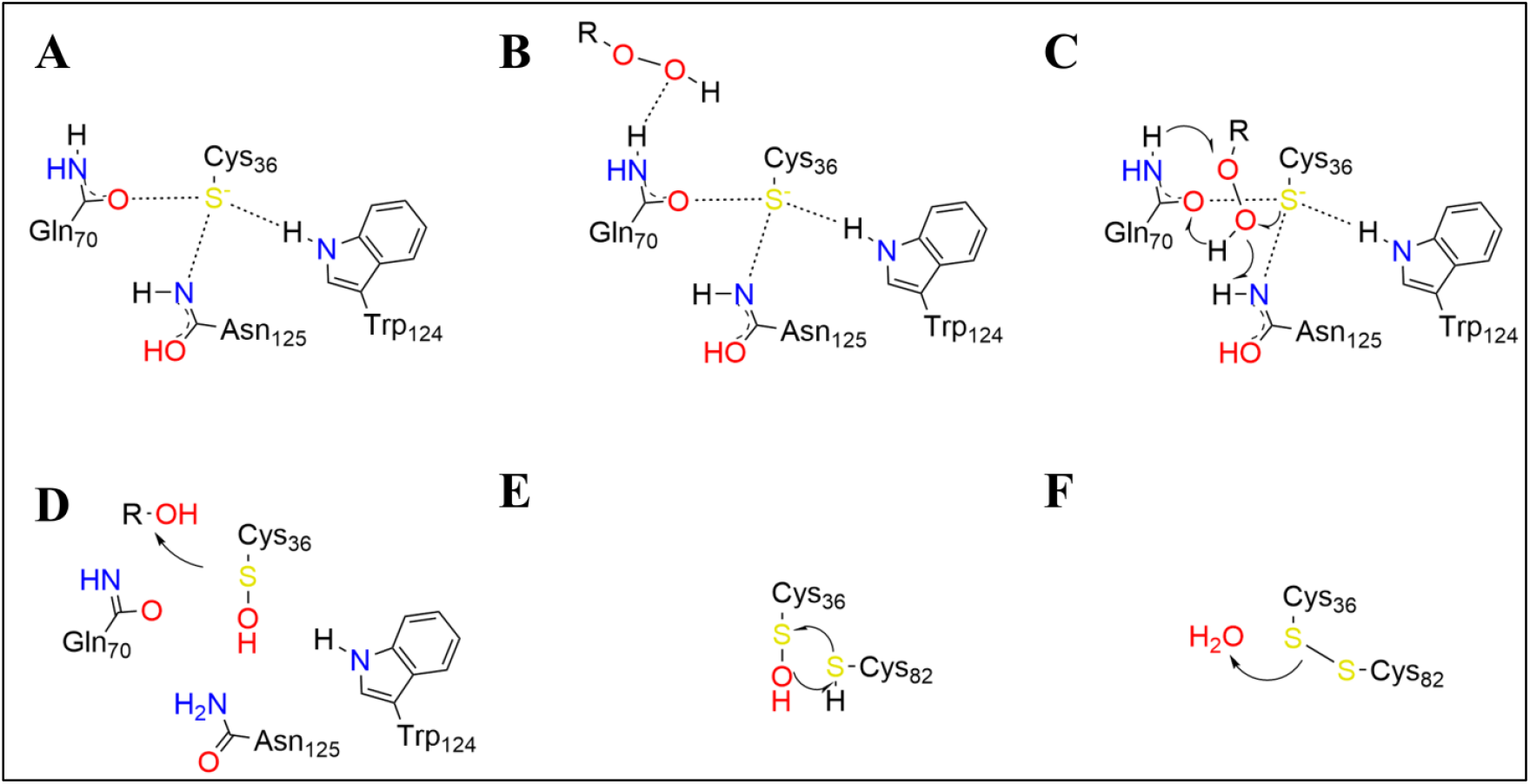
Possible catalytic mechanism of SaGpx. (8A) In the reduced state, the thiol group of C36 got converted into the nucleophilic thiolate anion form with the help of Q70, W124, and N125. (8B) The Q70, due to its surface exposed orientation, guide substrate hydroperoxide to the active site. (8C) The substrate hydroperoxide upon entering the active site, got stabilized by both Q70 and N125 which subsequently attached by thiolate anion of C36. (8D) The nucleophilic attack on substrate peroxide generates sulfenic acid intermediate with concomitant release of alcohol or water. (8E-F) Resolving cysteine, C82 then resolves the sulfenic acid intermediate into stable oxidized state with the release of one molecule of water.

Surface features of proteins play important roles from determining substrate specificity to make interaction with other proteins [^41^]. In spite of having sequence and structural similarities with other homologs, SaGpx exhibits distinct structural features that may lead to its functional adaptations in *S. aureus*. One such element is the presence of a positively charged cradle directly beneath the active site **(Fig. 7E)**. This cationic patch resembles the substrate binding cradle reported in human Gpx4 [^31^] **(Fig. 7A)** and the helminth isoform from *Schistosoma mansoni* [^29^], both of which utilize this region to stabilize bulky, negatively charged phospholipid hydroperoxides. In human Gpx4, this region is also reported to interact with DNA. The conservation of such a surface property in SaGpx strongly suggests that it too may act on complex lipid hydroperoxides, leading to broadening its substrate specificity beyond low molecular weight hydroperoxides. Interestingly, although poplar Gpx5 shares comparable sequence identity and global structural similarity with SaGpx, it lacks such a positively charged surface patch. Rather, poplar Gpx5 exhibits an overall distributed negative charge **(Fig. 7C, D)** that may facilitate different activity like providing protective role against heavy metal ion (for example cadmium) stress [^23^]. This difference highlights a possible divergence in substrate specificity where poplar Gpx5 is specialised for thioredoxin-dependent small molecule peroxide reduction along with to play a secondary role as heavy-metal sink [^23^], whereas SaGpx, akin to human Gpx4, extend its activity to membrane associated lipid peroxides. From a biological standpoint, the ability of SaGpx to detoxify phospholipid hydroperoxide could be advantageous for *S. aureus* in surviving oxidative bursts during host-pathogen interactions, protecting staphylococcal membrane from lipid peroxidation.

Beyond their classical role in hydroperoxide neutralization, glutathione peroxidases have also been reported to be involved in redox signalling processes. In *Saccharomyces cerevisiae*, the glutathione-dependent phospholipid peroxidase Hyr1 (also known as Gpx3) exerts this dual role. Hyr1 not only detoxifies peroxides but also acts as a redox sensor, transmitting oxidative signals to the transcription factor Yap1 through thiol-disulfide exchange mechanism [^45^]. This redox relay ultimately induces a wide array of antioxidant and stress responsive gene, providing the cell with adaptive protection against oxidative stress. SaGpx shares more than 50% amino acid sequence similarity and high structural resemblance of 0.54 Å (based on the alignment of 111 C_α_ positions) RMSD with Hyr1. This similarity highlights the possibility that SaGpx may also serve a similar function in redox signalling in *S. aureus*. This dual role would position SaGpx as a critical regulatory node, capable of both neutralizing oxidative threats and transducing redox information to downstream cellular processes.

Despite extensive crystallization attempts, wild type SaGpx failed to yield diffraction-quality crystals, likely due to oxidation-induced structural heterogeneity and intramolecular disulfide bond formation involving reactive cysteine residues. Crystal structure of wild type SaGpx would have provided the exact physiological three-dimensional arrangement of the catalytic cysteines and the overall structure of SaGpx. However, the C36S mutant successfully crystallized, reveals an active site architecture that is highly similar to that expected for the native enzyme, thereby offering a reliable structural surrogate for interpreting SaGpx mechanism and function. C36S SaGpx mutant structure provides the first high-resolution structure of a bacterial glutathione peroxidase. This cysteine to serine substitution eliminated redox-sensitive structural heterogeneity while preserving the overall protein architecture, enabling detailed structural characterization. Comparative structural analysis revealed significant homology with established Gpx enzymes, particularly with those acting as phospholipid hydroperoxidases and redox signalling mediators. These structural insights, combined with conserved functional domain strongly suggest that SaGpx may function beyond classical antioxidant roles, potentially serving as both a phospholipid hydroperoxidase and redox signalling transducer. Further experimental validation through structural explorations, biochemical assays and functional studies will be crucial to confirm these structure-based hypotheses and fully elucidate SaGpx’s physiological roles.

## Conclusion

This study presents the first structural characterization of a prokaryotic glutathione peroxidase homolog at high resolution (1.5 Å). The high-resolution crystal structure of *Staphylococcus aureus* glutathione peroxidase (C36S SaGpx mutant) reveals the presence of classical Gpx like conserved structural fold in SaGpx. Comparative sequential and structural analysis combined with site-directed mutagenesis and biochemical assays confirm the role of active site residues C36, Q70, W124, and N125, which collectively form the catalytic tetrad of this protein. Despite similarities with the classical mammalian selenocysteine-dependent Gpx enzymes, SaGpx was found to utilize two cysteine residues for the catalytic activity. Biochemical assays also confirm the thioredoxin-mediated catalytic regeneration of SaGpx following a similar catalytic mechanism as of atypical 2-cysteine peroxiredoxins. These finding supports the reclassification of SaGpx as thioredoxin-dependent glutathione peroxidase homolog. The monomeric SaGpx structure features a cationic patch near the active site, suggesting the possibility to function as a phospholipid hydroperoxidase which may impart comprehensive antioxidant protection during host-pathogen interactions. Also, comparative analysis with the glutathione-dependent phospholipid peroxidase Hyr1 from *Saccharomyces cerevisiae* suggests the plausible role of SaGpx in redox signalling processes. Collectively all this structure-centred hypothesis of different plausible physiological role of SaGpx has opened a vast area of future research, which with precise relevant experimental support, may enhance our knowledge in bacterial oxidative stress mitigation. Also, understanding these bacterial-specific anti-oxidative features may shed light on designing novel antibacterial therapeutics.

## Materials and Methods

### 1. Sequence analysis

The corresponding amino acid sequence of the putative Glutathione peroxidase from *Staphylococcus aureus* NCTC8325 strain was retrieved from KEGG organism database [^20^]. With this amino acid sequence, NCBI protein BLAST [^46^] search was performed against PDB database [^47^] to find out Gpx homologs showing more than 40% sequence similarity with SaGpx. The amino acid sequence of SaGpx was then subjected to sequence alignment with the obtained Gpx homologs from different organism upon BLAST search. Apart from BLAST result, amino acid sequences of human Gpx’s were considered for comparative analysis. The sequence alignment was performed with ClustalW [^48^]. Upon performing alignment, the alignment data was analysed through Espript [^49^].

### 2. Cloning, over-expression and purification

The corresponding ORFs of the target constructs i.e., SaGpx (SAOUHSC_01282), SaTrx1 (SAOUHSC_01100), SaTrx2 (SAOUHSC_00834) and SaTR (SAOUHSC_00785) from *Staphylococcus aureus* RN4220 strain (descendant strain of *Staphylococcus aureus* strain NCTC8325-4) were cloned into pET28 (for SaGpx) and pQE30 (for SaTrx1, SaTrx2 and SaTR) plasmid vector. The recombinant plasmid adds a hexa-histidine (6xHis) tag to the N-terminus of the protein which is then over-expressed in *E. coli* BL21 DE3, *E. coli* M15 or *E. coli* SG13009 expression host. Bacterial cells were grown on LB culture media at 37º C up to an O.D of ∼0.6 at 600 nm wavelength. Expression was induced with 100 µM of Isopropyl β-D-1-thiogalactopyranoside (IPTG) and cells were harvested after 4 hours of incubation at 37º C. Harvested cells were resuspended in lysis buffer (10 mM Tris pH 8.0, 300 mM NaCl and 10 mM Imidazole) and lysed by sonication. The sonicated sample was then centrifuged and the supernatant part containing the soluble target protein fraction was collected and loaded on to a Ni-NTA column (Cytiva) equilibrated with the binding buffer (10 mM Tris pH 8.0, 300 mM NaCl and 10 mM Imidazole). The target protein was eluted with different gradient of Imidazole (10 mM to 300 mM) in elution buffer (10 mM Tris pH 8.0, 300 mM NaCl and varying concentration of Imidazole). The eluted fractions from each different gradient of Imidazole were then collected and ran on 12% or 15% SDS-PAGE to identify the eluted fraction containing the target protein. The selected fraction is then taken for further purification and loaded on to a Superdex 75 pg size exclusion chromatography column (Cytiva) already equilibrated with gel filtration buffer (for SaGpx, 20 mM Tris pH 8.0, 150 mM NaCl and 5 mM DTT, whereas for SaTrx1, SaTrx2 and SaTR, 20 mM Tris pH 8.0, 50 mM NaCl and 5 mM DTT was used). The fraction showing single peak at λ_Max._ = 280 nm on the chromatogram was collected and stored at −80º C with final 5% glycerol as cryo-protecting agent. This stored protein was used later in corresponding enzyme assays. Site-directed mutagenesis of Wild type SaGpx was performed at positions 36^th^, 64^th^, 70^th^, 82^nd^, 124^th^, 125^th^, (64^th^ and 82^nd^) to get C36S, C64S, C82S, Q70A, W124A, N125A and (C64S+C82S) mutants. The site-directed mutagenesis was performed using QuikChange II Site-Directed Mutagenesis Kit (Agilent) as per the given protocol. The mutants were cloned in pET28 plasmid vector and expressed in *E. coli* BL21 DE3 expression host. For expression and purification of different SaGpx mutants, same protocol was followed as of wild type SaGpx.

### 3. Enzyme activity assay of wild type SaGpx and comparative analysis with its different mutants

Wild type SaGpx and C36S, C64S, C82S, Q70A, W124A, N125A and (C64S+C82S) mutants of SaGpx proteins were homogenously purified in reducing conditions. Before enzymatic analysis, each protein was buffer exchanged with the corresponding gel filtration buffer to remove DTT and Glycerol by passing through PD10 desalting column (Cytiva). All the UV-Vis spectroscopic readings were recorded using SHIMADZU UV-1900 UV-Vis Spectrophotometer.

#### 3.1. Peroxidase activity assessment of SaGpx and its various site-directed mutants

Peroxidase activity was measured with the help of Ellman’s reagent or 5,5’-dithio-bis-(2-nitrobenzoic acid), also known as DTNB [^50^]. This colorimetric method is based on the detection of free thiol group of SaGpx and its mutant proteins in the presence or absence of substrate t-BOOH. Assay was performed in reaction buffer containing 20 mM Tris pH 8.0 and 150 mM NaCl. Stock solution (10 mM) of DTNB was prepared in 100% DMSO. Protein concentration was kept constant at 20 µM with varying t-BOOH concentrations (0, 0.2, 0.5, 1, 2, 5, 10, 20, 30, and 50 µM). Total reaction volume was of 250 µL. The reaction was started upon addition of t-BOOH with SaGpx or its mutants in buffered solution. Then the assay mixture was incubated for 2 minutes at room temperature. After incubation, the reaction was stopped by adding 250 µL 10% SDS. Then 25 µL of 10 mM DTNB solution was added and incubated for another 30 minutes. Finally, absorbance at 412 nm was recorded. All the experiments were performed in triplicate.

#### 3.2. Comparative substrate preference analysis of SaGpx

Coupled assay as previously reported methodology [^51^] was performed with some modifications to find out the favoured peroxide substrate of SaGpx. Three different hydroperoxides namely hydrogen peroxide (H_2_O_2_), tert-Butyl hydroperoxide (t-BOOH) and cumene hydroperoxide were used in this assay. The reaction mixture contains NADPH, SaTR, SaTrx1, SaGpx, and hydroperoxide. In each case, the concentration of NADPH, SaTR, SaTrx1, and SaGpx are 0.2mM, 1.5μM, 2μM, and 5μM respectively with varying hydroperoxide (H_2_O_2_/t-BOOH/cumene hydroperoxide) concentrations (0µM, 10 µM, 15 µM, 20 µM, 25 µM, 50 µM, 100 µM, 150 µM, 200 µM, 500 µM, 750 µM and 1000 µM). The reaction buffer contains 20 mM Tris pH 8.0 and 150 mM NaCl. First, appropriate volume of SaTR, SaTrx1, and SaGpx were mixed and incubated for 1 hour at 4 °C. Then in a microcentrifuge tube Milli-Q water, appropriate volume of 10x reaction buffer, NADPH and pre-incubated protein mixture was added sequentially. Then the reaction was started by adding varying concentration of different hydroperoxide substrate in each case. After adding hydroperoxide, the absorbance at 340nm was taken immediately and monitored for 1 minute. In each case the reaction mixture without SaGpx was used as a control. All the reactions were performed in triplicate. Absorbance values were plotted in Origin software and different kinetic parameters like K_m_, V_max_, K_cat_ and catalytic efficiency (K_cat_/K_m_) were calculated for probing SaGpx’s substrate specificity.

#### 3.3. Cognate electron donor specificity analysis of SaGpx

After probing the preferred substrate hydroperoxide for SaGpx, once again the coupled assay of the SaGpx with SaTrx1 or SaTrx2 was performed to find out which one is the plausible cognate electron donor protein. The reaction mixture contains NADPH, SaTR, SaTrx1/SaTrx2, SaGpx, and cumene hydroperoxide. In each case, the concentration of NADPH, SaTR, SaGpx and cumene hydroperoxide were kept 0.2 mM, 1.5 μM, 5 μM, and 100 μM respectively with varying SaTrx1/SaTrx2 concentrations (0 µM, 1 µM, 2 µM, 3 µM, 5 µM, 7.5 µM, 10 µM, 12.5 µM, 15 µM, and 20 µM). The reaction buffer contains 20 mM Tris pH 8.0 and 150 mM NaCl. First, appropriate volume of SaTR, SaTrx1/SaTrx2, and SaGpx were mixed and incubated for 1 hour at 4 °C. Then in a microcentrifuge tube Milli-Q water, appropriate volume of 10x buffer, NADPH and pre-incubated protein mixture were added sequentially. Then the reaction was started by adding cumene hydroperoxide. After adding cumene hydroperoxide, the absorbance at 340nm was taken immediately and monitored for 1 minute. In each case the reaction mixture without SaTrx1 or SaTrx2 was used as a control. All the reactions were performed in triplicate. Kinetic parameters were calculated for probing possible cognate electron donor specificity for SaGpx.

#### 3.4. Comparison of catalytic activity between Wild type SaGpx and its mutants

To elucidate the role of each of the highly conserved active site residues of SaGpx in the electron relay from NADPH to the hydroperoxide substrate, coupled assay was performed (as described in 3.2. section with cumene hydroperoxide as substrate) with each of the created point mutants of SaGpx, namely C36S, C64S, C82S, Q70A, W124A, N125A. With these coupled assays, different kinetic parameters were calculated (K_m_, V_max_, K_cat_ and the catalytic efficiency, K_cat_/K_m_) for each of the mutants. These kinetic values were compared with the wild type SaGpx and the role of each of the active site amino acid residues in the catalytic activity of SaGpx was determined.

#### 3.5. Oxidation dependent conformational change analysis of SaGpx using CD spectroscopy and Fluorescence spectroscopy

Wild type SaGpx was used in both experiments. Protein and t-BOOH concentration was kept at 5 µM and 50 µM respectively. The reaction buffer composed of 20 mM Tris pH 8.0, 150 mM NaCl. For Circular Dichroism (CD) spectroscopy analysis, a spectral scan from 190 nm to 350 nm was recorded using JASCO J-815 CD Spectrometer. For Fluorescence spectroscopy, JASCO FP-8300 Spectrofluorometer was used to record emission spectra from 310nm to 700nm while keeping excitation maxima at 295 nm.

#### 3.6. Detection of cysteine-sulfenic acid intermediate in SaGpx

Formation of sulfenic acid was probed by using 4-Chloro-7-nitrobenzofurazan or NBD chloride (NBD-Cl) [^36^]. C64S+C82S mutant protein was used in this experiment. Reduced protein was incubated with t-BOOH in 1:1 molar ratio at room temperature to get all thiols converted into sulfenic acid form. Reduced protein without any treatment serves as control. Subsequently, both reduced and oxidized proteins were incubated with NBD-Cl at a 10-fold molar excess relative to protein concentration for 30 minutes at room temperature. After incubation both the protein samples were placed in a 10 kDa concentrator and excess NBD-Cl was removed by repeated cycles of concentration and dilution with reaction buffer (20 mM Tris pH 8.0, 150 mM NaCl). After removing excess NBD-Cl, the concentration of the washed protein samples was adjusted to ∼50 µM by taking absorbance at 280 nm. Finally, a spectral scan from 200 nm to 500 nm was recorded for both the protein sample and compared at 343 nm and 420 nm, the characteristic peaks for NBD-Cl adduct with oxidized and reduced proteins.

### 4. Crystallization of C36S SaGpx mutant

For Crystallization, the salt concentration of gel filtration buffer was increased from 150 mM to 250 mM with 20 mM Tris pH 8.0 and 5 mM DTT. The high salt concentration is required to prevent the observed precipitation of concentrated protein (around 20 mg/ml) required for crystallization. The homogenously purified protein sample was used for crystallization study. Crystallization conditions for C36S mutant were screened extensively with Crystal Screen (CS) and Index Screen (Hampton Research) crystallization sparse matrix solutions. The crystals were grown at room temperature (∼298 K). Rough screening was done using sitting drop method on 96 well Intelli-Plate (Art Robbins Instruments) by mixing equal volume of protein sample (22 mg/ml) with crystallization sparse matrix solutions. Among several conditions, G4 condition (1.6 M sodium citrate tribasic dihydrate pH 6.5) from CS shows promising crystal features upon initial screening trials. This condition was then taken further for fine crystallization screening. Fine crystallization screening was performed on 24-well Linbro plates (Hi-Media) by using hanging drop vapour diffusion method. Drops were casted on siliconized glass coverslip by mixing 2 µL of concentrated protein sample (22mg/mL) with 2 µL of reservoir solution of fine screening conditions. Crystals appeared within 48-96 hours of incubation at room temperature (∼298 K). After necessary optimization of pH, salt (Sodium citrate tribasic dihydrate) concentration, diffraction quality crystals were obtained. All the crystals show plate like morphology originating from a central hub **(Fig. S8A)**. Crystals were tweaked from the central hub and snap cooled in liquid nitrogen. As the crystals appeared in high salt concentration, no additional cryo-protectants were used during diffracting the crystal with X-ray in cryogenic conditions (temperature, 100K). **(Fig. S8B)**. Finally, crystals appeared from 1.2 M sodium citrate tribasic dihydrate pH 6.0 condition were diffracted at high resolution.

### 5. Data collection and Structure solution

Crystals were diffracted with single wavelength (λ _X-ray_ = 0.978930 Å) X-ray beam at PX-BL21 beamline facility at Indus-2, RRCAT, Indore, India [^52^]. Diffraction datasets were collected by using MARCCD 225 (Rayonix) detector with continuous flow of liquid nitrogen stream at 100K temperature. A total of 360 frames were collected with 1º oscillation by keeping the detector at 200 nm distance. The diffraction dataset were reduced, indexed and integrated with XDS software package [^53^]. Then the point group determination and scaling of the reduced dataset was performed by using Pointless [^54^] and SCALA [^54^] respectively from CCP4 suite [^55^]. The number of protein molecules in the asymmetric unit was estimated using the Matthews coefficient [^56^]. Phasing was performed by Molecular replacement using MOLREP [^57^] or Phaser MR of CCP4 suite. The crystal structure of the poplar glutathione peroxidase 5 in the reduced form (PDB Id: 2P5Q) was used as template in molecular replacement [^23^]. The model obtained after phasing was subjected to rigid body refinement followed by restrain body refinement using Refmac5 [^58^] of CCP4 suite or Phenix [^59^] and visualization was performed on Coot [^60^]. Iterative cycles of model building and refinement were performed till synchronised agreement between the structure model and experimental data is obtained. Finally, the model structure was stereochemically validated and submitted to PDB database [^47^] with the PDB ID: 9KZM.

## Abbreviations

SaGpx: staphylococcal glutathione peroxidase
SaTrx1: staphylococcal thioredoxin 1
SaTrx2: staphylococcal thioredoxin 2
ROS: reactive oxygen species
Gpx: glutathione peroxidase
Trx: thioredoxin
TR: thioredoxin reductase
GR: glutathione reductase
GSH: reduced glutathione
GSSG: oxidized glutathione
Prx: peroxiredoxin
NADPH: reduced nicotinamide adenine dinucleotide phosphate
NADP: oxidized nicotinamide adenine dinucleotide phosphate
DNA: deoxyribonucleic acid
DTNB: 5,5’-dithio-bis-(2-nitrobenzoic acid)
TNB: 2-nitro-5-thiobenzoic acid
DTT: dithiothreitol.

## Acknowledgements

The study is supported by grants from Start-up Research Grant (SRG) funded by Science and Engineering Research Board (SERB), Department of Science and Technology, Government of India (Sanction no. SRG/2020/001353), SEED grant supported by IIT Jodhpur (Sanction no. I/SEED/SUB/20200005) and AYURTECH facility at IIT Jodhpur, Ministry of AYUSH, Government of India (Sanction no. S-12011/12/2021-SCHEME). Authors including S.M. and M.S. expresses special thanks to Ministry of Education, Government of India for providing research fellowships to carry out this work. Authors are thankful to IIT Jodhpur for providing experimental facility and necessary infrastructure for carrying out this work. Authors are also thankful to PX-BL21 beamline (BARC) at Indus-2, RRCAT, Indore, India for protein crystal diffraction and subsequent data collection facility.

## Author Contributions

**SM:** Investigation, Methodology, Data curation, Formal analysis, Writing –original draft **MS:** Data curation **SB:** Conceptualization, Project administration, Supervision, Writing – review and editing, Resources and Funding acquisition

## References

(1) Lennicke, C.; Cochemé, H. M. Redox Metabolism: ROS as Specific Molecular Regulators of Cell Signaling and Function. Mol. Cell 2021, 81 (18), 3691–3707. 10.1016/j.molcel.2021.08.018.

(2) Murphy, M. P. How Mitochondria Produce Reactive Oxygen Species. Biochem. J. 2009, 417 (1), 1–13. 10.1042/BJ20081386.

(3) Zorov, D. B.; Juhaszova, M.; Sollott, S. J. Mitochondrial Reactive Oxygen Species (ROS) and ROS-Induced ROS Release. Physiol. Rev. 2014, 94 (3), 909–950. 10.1152/physrev.00026.2013.

(4) Forrester, S. J.; Kikuchi, D. S.; Hernandes, M. S.; Xu, Q.; Griendling, K. K. Reactive Oxygen Species in Metabolic and Inflammatory Signaling. Circ. Res. 2018, 122 (6), 877–902. 10.1161/CIRCRESAHA.117.311401.

(5) Nguyen, G. T.; Green, E. R.; Mecsas, J. Neutrophils to the ROScue: Mechanisms of NADPH Oxidase Activation and Bacterial Resistance. Front. Cell. Infect. Microbiol. 2017, 7, 373. 10.3389/fcimb.2017.00373.

(6) Phaniendra, A.; Jestadi, D. B.; Periyasamy, L. Free Radicals: Properties, Sources, Targets, and Their Implication in Various Diseases. Indian J. Clin. Biochem. 2015, 30 (1), 11–26. 10.1007/s12291-014-0446-0.

(7) Jomova, K.; Alomar, S. Y.; Alwasel, S. H.; Nepovimova, E.; Kuca, K.; Valko, M. Several Lines of Antioxidant Defense against Oxidative Stress: Antioxidant Enzymes, Nanomaterials with Multiple Enzyme-Mimicking Activities, and Low-Molecular-Weight Antioxidants. Arch. Toxicol. 2024, 98 (5), 1323–1367. 10.1007/s00204-024-03696-4.

(8) Ighodaro, O. M.; Akinloye, O. A. First Line Defence Antioxidants-Superoxide Dismutase (SOD), Catalase (CAT) and Glutathione Peroxidase (GPX): Their Fundamental Role in the Entire Antioxidant Defence Grid. Alex. J. Med. 2018, 54 (4), 287–293. 10.1016/j.ajme.2017.09.001.

(9) Mishra, S.; Imlay, J. Why Do Bacteria Use so Many Enzymes to Scavenge Hydrogen Peroxide? Arch. Biochem. Biophys. 2012, 525 (2), 145–160. 10.1016/j.abb.2012.04.014.

(10) Flohé, L.; Jaeger, T.; Pilawa, S.; Sztajer, H. Thiol-Dependent Peroxidases Care Little about Homology-Based Assignments of Function. Redox Rep. 2003, 8 (5), 256–264. 10.1179/135100003225002862.

(11) Brigelius-Flohé, R.; Maiorino, M. Glutathione Peroxidases. Biochim. Biophys. Acta BBA - Gen. Subj. 2013, 1830 (5), 3289– 3303. 10.1016/j.bbagen.2012.11.020.

(12) Lu, J.; Holmgren, A. The Thioredoxin Antioxidant System. Free Radic. Biol. Med. 2014, 66, 75–87. 10.1016/j.freeradbiomed.2013.07.036.

(13) Pei, J.; Pan, X.; Wei, G.; Hua, Y. Research Progress of Glutathione Peroxidase Family (GPX) in Redoxidation. Front. Pharmacol. 2023, 14, 1147414. 10.3389/fphar.2023.1147414.

(14) Minard, K. I.; Jennings, G. T.; Loftus, T. M.; Xuan, D.; McAlister-Henn, L. Sources of NADPH and Expression of Mammalian NADP+-Specific Isocitrate Dehydrogenases in Saccharomyces Cerevisiae. J. Biol. Chem. 1998, 273 (47), 31486–31493. 10.1074/jbc.273.47.31486.

(15) Brenot, A.; King, K. Y.; Janowiak, B.; Griffith, O.; Caparon, M. G. Contribution of Glutathione Peroxidase to the Virulence of Streptococcus Pyogenes. Infect. Immun. 2004, 72 (1), 408–413. 10.1128/IAI.72.1.408-413.2004.

(16) Zhang, Y.; Guo, Q.; Fang, X.; Yuan, M.; Hu, W.; Liang, X.; Liu, J.; Yang, Y.; Fang, C. Destroying Glutathione Peroxidase Improves the Oxidative Stress Resistance and Pathogenicity of Listeria Monocytogenes. Front. Microbiol. 2023, 14, 1122623. 10.3389/fmicb.2023.1122623.

(17) Gaupp, R.; Ledala, N.; Somerville, G. A. Staphylococcal Response to Oxidative Stress. Front. Cell. Infect. Microbiol. 2012, 2. 10.3389/fcimb.2012.00033.

(18) Pidwill, G. R.; Gibson, J. F.; Cole, J.; Renshaw, S. A.; Foster, S. J. The Role of Macrophages in Staphylococcus Aureus Infection. Front. Immunol. 2021, 11, 620339. 10.3389/fimmu.2020.620339.

(19) Voyich, J. M.; Braughton, K. R.; Sturdevant, D. E.; Whitney, A. R.; Saïd-Salim, B.; Porcella, S. F.; Long, R. D.; Dorward, D. W.; Gardner, D. J.; Kreiswirth, B. N.; Musser, J. M.; DeLeo, F. R. Insights into Mechanisms Used by Staphylococcus Aureus to Avoid Destruction by Human Neutrophils. J. Immunol. 2005, 175 (6), 3907–3919. 10.4049/jimmunol.175.6.3907.

(20) Kanehisa, M. KEGG: Kyoto Encyclopedia of Genes and Genomes. Nucleic Acids Res. 2000, 28 (1), 27–30. 10.1093/nar/28.1.27.

(21) Uziel, O.; Borovok, I.; Schreiber, R.; Cohen, G.; Aharonowitz, Y. Transcriptional Regulation of the Staphylococcus Aureus Thioredoxin and Thioredoxin Reductase Genes in Response to Oxygen and Disulfide Stress. J. Bacteriol. 2004, 186 (2), 326– 334. 10.1128/JB.186.2.326-334.2004.

(22) Lensmire, J. M.; Wischer, M. R.; Kraemer-Zimpel, C.; Kies, P. J.; Sosinski, L.; Ensink, E.; Dodson, J. P.; Shook, J. C.; Delekta, P. C.; Cooper, C. C.; Havlichek, D. H.; Mulks, M. H.; Lunt, S. Y.; Ravi, J.; Hammer, N. D. The Glutathione Import System Satisfies the Staphylococcus Aureus Nutrient Sulfur Requirement and Promotes Interspecies Competition. PLOS Genet. 2023, 19 (7), e1010834. 10.1371/journal.pgen.1010834.

(23) Koh, C. S.; Didierjean, C.; Navrot, N.; Panjikar, S.; Mulliert, G.; Rouhier, N.; Jacquot, J.-P.; Aubry, A.; Shawkataly, O.; Corbier, C. Crystal Structures of a Poplar Thioredoxin Peroxidase That Exhibits the Structure of Glutathione Peroxidases: Insights into Redox-Driven Conformational Changes. J. Mol. Biol. 2007, 370 (3), 512–529. 10.1016/j.jmb.2007.04.031.

(24) Zhang, W.; He, Y.; Yang, Z.; Yu, J.; Chen, Y.; Zhou, C. Crystal Structure of Glutathione‐dependent Phospholipid Peroxidase Hyr1 from the Yeast Saccharomyces Cerevisiae. Proteins Struct. Funct. Bioinforma. 2008, 73 (4), 1058–1062. 10.1002/prot.22220.

(25) Maiorino, M.; Ursini, F.; Bosello, V.; Toppo, S.; Tosatto, S. C. E.; Mauri, P.; Becker, K.; Roveri, A.; Bulato, C.; Benazzi, L.; De Palma, A.; Flohé, L. The Thioredoxin Specificity of Drosophila GPx: A Paradigm for a Peroxiredoxin-like Mechanism of Many Glutathione Peroxidases. J. Mol. Biol. 2007, 365 (4), 1033–1046. 10.1016/j.jmb.2006.10.033.

(26) Wood, Z. A.; Schröder, E.; Robin Harris, J.; Poole, L. B. Structure, Mechanism and Regulation of Peroxiredoxins. Trends Biochem. Sci. 2003, 28 (1), 32–40. 10.1016/S0968-0004(02)00003-8.

(27) Adriani, P. P.; de Paiva, F. C. R.; de Oliveira, G. S.; Leite, A. C.; Sanches, A. S.; Lopes, A. R.; Dias, M. V. B.; Chambergo, F. S. Structural and Functional Characterization of the Glutathione Peroxidase-like Thioredoxin Peroxidase from the Fungus Trichoderma Reesei. Int. J. Biol. Macromol. 2021, 167, 93–100. 10.1016/j.ijbiomac.2020.11.179.

(28) Patel, S.; Hussain, S.; Harris, R.; Sardiwal, S.; Kelly, J. M.; Wilkinson, S. R.; Driscoll, P. C.; Djordjevic, S. Structural Insights into the Catalytic Mechanism of Trypanosoma Cruzi GPXI (Glutathione Peroxidase-like Enzyme I). Biochem. J. 2010, 425 (3), 513–522. 10.1042/BJ20091167.

(29) Dimastrogiovanni, D.; Anselmi, M.; Miele, A. E.; Boumis, G.; Petersson, L.; Angelucci, F.; Nola, A. D.; Brunori, M.; Bellelli, Combining Crystallography and Molecular Dynamics: The Case of Schistosoma Mansoni Phospholipid Glutathione Peroxidase. Proteins Struct. Funct. Bioinforma. 2010, 78 (2), 259–270. 10.1002/prot.22536.

(30) Alphey, M. S.; König, J.; Fairlamb, A. H. Structural and Mechanistic Insights into Type II Trypanosomatid Tryparedoxin-Dependent Peroxidases. Biochem. J. 2008, 414 (3), 375–381. 10.1042/BJ20080889.

(31) Scheerer, P.; Borchert, A.; Krauss, N.; Wessner, H.; Gerth, C.; Höhne, W.; Kuhn, H. Structural Basis for Catalytic Activity and Enzyme Polymerization of Phospholipid Hydroperoxide Glutathione Peroxidase-4 (GPx4)^;^. Biochemistry 2007, 46 (31), 9041– 9049. 10.1021/bi700840d.

(32) Savelli, B.; Li, Q.; Webber, M.; Jemmat, A. M.; Robitaille, A.; Zamocky, M.; Mathé, C.; Dunand, C. RedoxiBase: A Database for ROS Homeostasis Regulated Proteins. Redox Biol. 2019, 26, 101247. 10.1016/j.redox.2019.101247.

(33) Chen, Y.; Barkley, M. D. Toward Understanding Tryptophan Fluorescence in Proteins. Biochemistry 1998, 37 (28), 9976–9982. 10.1021/bi980274n.

(34) Paulsen, C. E.; Carroll, K. S. Cysteine-Mediated Redox Signaling: Chemistry, Biology, and Tools for Discovery. Chem. Rev. 2013, 113 (7), 4633–4679. 10.1021/cr300163e.

(35) Baker, L. M. S.; Poole, L. B. Catalytic Mechanism of Thiol Peroxidase from Escherichia Coli. J. Biol. Chem. 2003, 278 (11), 9203–9211. 10.1074/jbc.M209888200.

(36) Poole, L. B. Measurement of Protein Sulfenic Acid Content. Curr. Protoc. Toxicol. 2008, 38 (1). 10.1002/0471140856.tx1702s38.

(37) Martin, J. L. Thioredoxin —a Fold for All Reasons. Structure 1995, 3 (3), 245–250. 10.1016/S0969-2126(01)00154-X.

(38) Webb, B.; Sali, A. Comparative Protein Structure Modeling Using MODELLER. Curr. Protoc. Bioinforma. 2016, 54 (1). 10.1002/cpbi.3.

(39) Holm, L.; Laiho, A.; Törönen, P.; Salgado, M. DALI Shines a Light on Remote Homologs: One Hundred Discoveries. Protein Sci. 2023, 32 (1), e4519. 10.1002/pro.4519.

(40) Nguyen, V. D.; Saaranen, M. J.; Karala, A.-R.; Lappi, A.-K.; Wang, L.; Raykhel, I. B.; Alanen, H. I.; Salo, K. E. H.; Wang, C.; Ruddock, L. W. Two Endoplasmic Reticulum PDI Peroxidases Increase the Efficiency of the Use of Peroxide during Disulfide Bond Formation. J. Mol. Biol. 2011, 406 (3), 503–515. 10.1016/j.jmb.2010.12.039.

(41) Labrecque, C. L.; Fuglestad, B. Electrostatic Drivers of GPx4 Interactions with Membrane, Lipids, and DNA. Biochemistry 2021, 60 (37), 2761–2772. 10.1021/acs.biochem.1c00492.

(42) Flohé, L.; Toppo, S.; Cozza, G.; Ursini, F. A Comparison of Thiol Peroxidase Mechanisms. Antioxid. Redox Signal. 2011, 15 (3), 763–780. 10.1089/ars.2010.3397.

(43) Wood, Z. A.; Schröder, E.; Robin Harris, J.; Poole, L. B. Structure, Mechanism and Regulation of Peroxiredoxins. Trends Biochem. Sci. 2003, 28 (1), 32–40. 10.1016/S0968-0004(02)00003-8.

(44) Tosatto, S. C. E.; Bosello, V.; Fogolari, F.; Mauri, P.; Roveri, A.; Toppo, S.; Flohé, L.; Ursini, F.; Maiorino, M. The Catalytic Site of Glutathione Peroxidases. Antioxid. Redox Signal. 2008, 10 (9), 1515–1526. 10.1089/ars.2008.2055.

(45) Delaunay, A.; Pflieger, D.; Barrault, M.-B.; Vinh, J.; Toledano, M. B. A Thiol Peroxidase Is an H2O2 Receptor and Redox-Transducer in Gene Activation. Cell 2002, 111 (4), 471–481. 10.1016/S0092-8674(02)01048-6.

(46) Altschul, S. F.; Gish, W.; Miller, W.; Myers, E. W.; Lipman, D. J. Basic Local Alignment Search Tool. J. Mol. Biol. 1990, 215 (3), 403–410. 10.1016/S0022-2836(05)80360-2.

(47) Berman, H. M. The Protein Data Bank. Nucleic Acids Res. 2000, 28 (1), 235–242. 10.1093/nar/28.1.235.

(48) Thompson, J. D.; Higgins, D. G.; Gibson, T. J. CLUSTAL W: Improving the Sensitivity of Progressive Multiple Sequence Alignment through Sequence Weighting, Position-Specific Gap Penalties and Weight Matrix Choice. Nucleic Acids Res. 1994, 22 (22), 4673–4680. 10.1093/nar/22.22.4673.

(49) Gouet, P.; Courcelle, E.; Stuart, D. I.; M√©toz, F. ESPript: Analysis of Multiple Sequence Alignments in PostScript. Bioinformatics 1999, 15 (4), 305–308. 10.1093/bioinformatics/15.4.305.

(50) Ellman, G. L. Tissue Sulfhydryl Groups. Arch. Biochem. Biophys. 1959, 82 (1), 70–77. 10.1016/0003-9861(59)90090-6.

(51) Delaunay, A.; Pflieger, D.; Barrault, M.-B.; Vinh, J.; Toledano, M. B. A Thiol Peroxidase Is an H2O2 Receptor and Redox-Transducer in Gene Activation. Cell 2002, 111 (4), 471–481. 10.1016/S0092-8674(02)01048-6.

(52) Kumar, A.; Ghosh, B.; Poswal, H. K.; Pandey, K. K.; Jagannath Hosur, M.V.; Dwivedi, A.; Makde, R. D.; Sharma, S. M. Protein Crystallography Beamline (PX-BL21) at Indus-2 Synchrotron. J. Synchrotron Radiat. 2016, 23 (2), 629–634. 10.1107/S160057751600076X.

(53) Kabsch, W. XDS. Acta Crystallogr. D Biol. Crystallogr. 2010, 66 (2), 125–132. 10.1107/S0907444909047337.

(54) Evans, P. R. An Introduction to Data Reduction: Space-Group Determination, Scaling and Intensity Statistics. Acta Crystallogr. D Biol. Crystallogr. 2011, 67 (4), 282–292. 10.1107/S090744491003982X.

(55) Potterton, E.; Briggs, P.; Turkenburg, M.; Dodson, E. A Graphical User Interface to the CCP 4 Program Suite. Acta Crystallogr. D Biol. Crystallogr. 2003, 59 (7), 1131–1137. 10.1107/S0907444903008126.

(56) Kantardjieff, K. A.; Rupp, B. Matthews Coefficient Probabilities: Improved Estimates for Unit Cell Contents of Proteins, DNA, and Protein–Nucleic Acid Complex Crystals. Protein Sci. 2003, 12 (9), 1865–1871. 10.1110/ps.0350503.

(57) Vagin, A.; Teplyakov, A. MOLREP: An Automated Program for Molecular Replacement. J. Appl. Crystallogr. 1997, 30 (6), 1022–1025. 10.1107/S0021889897006766.

(58) Murshudov, G. N.; Skubák, P.; Lebedev, A. A.; Pannu, N. S.; Steiner, R. A.; Nicholls, R. A.; Winn, M. D.; Long, F.; Vagin, A. REFMAC 5 for the Refinement of Macromolecular Crystal Structures. Acta Crystallogr. D Biol. Crystallogr. 2011, 67 (4), 355–367. 10.1107/S0907444911001314.

(59) Liebschner, D.; Afonine, P. V.; Baker, M. L.; Bunkóczi, G.; Chen, V. B.; Croll, T. I.; Hintze, B.; Hung, L.-W.; Jain, S.; McCoy, J.; Moriarty, N. W.; Oeffner, R. D.; Poon, B. K.; Prisant, M. G.; Read, R. J.; Richardson, J. S.; Richardson, D. C.; Sammito, M. D.; Sobolev, O. V.; Stockwell, D. H.; Terwilliger, T. C.; Urzhumtsev, A. G.; Videau, L. L.; Williams, C. J.; Adams, P. D. Macromolecular Structure Determination Using X-Rays, Neutrons and Electrons: Recent Developments in Phenix. Acta Crystallogr. Sect. Struct. Biol. 2019, 75 (10), 861–877. 10.1107/S2059798319011471.

(60) Emsley, P.; Lohkamp, B.; Scott, W. G.; Cowtan, K. Features and Development of Coot. Acta Crystallogr. D Biol. Crystallogr. 2010, 66 (4), 486–501. 10.1107/S0907444910007493.

