## Supplementary Data for "Deciphering the Structure and Mechanism of SaGpx: A Non-Canonical Glutathione Peroxidase from *Staphylococcus aureus*"

### 2. Materials and Methods:

#### 2.1. Fine chemicals, reagents and enzymes:

All the fine chemicals and reagents needed for the experiments described in the study were of analytical grade and purchased from SRL (India), Qualigens (India), HiMedia (India) and Sigma (USA). The enzymes used in cloning of the target constructs were purchased from NEB (USA) and Promega (USA). All the matrix used in Ni-NTA column and size-exclusion column were purchased from Cytiva (USA). All the crystallization sparse matrix screens and associated products used in crystallization were purchased from Hampton Research (USA).

### 3. Results:

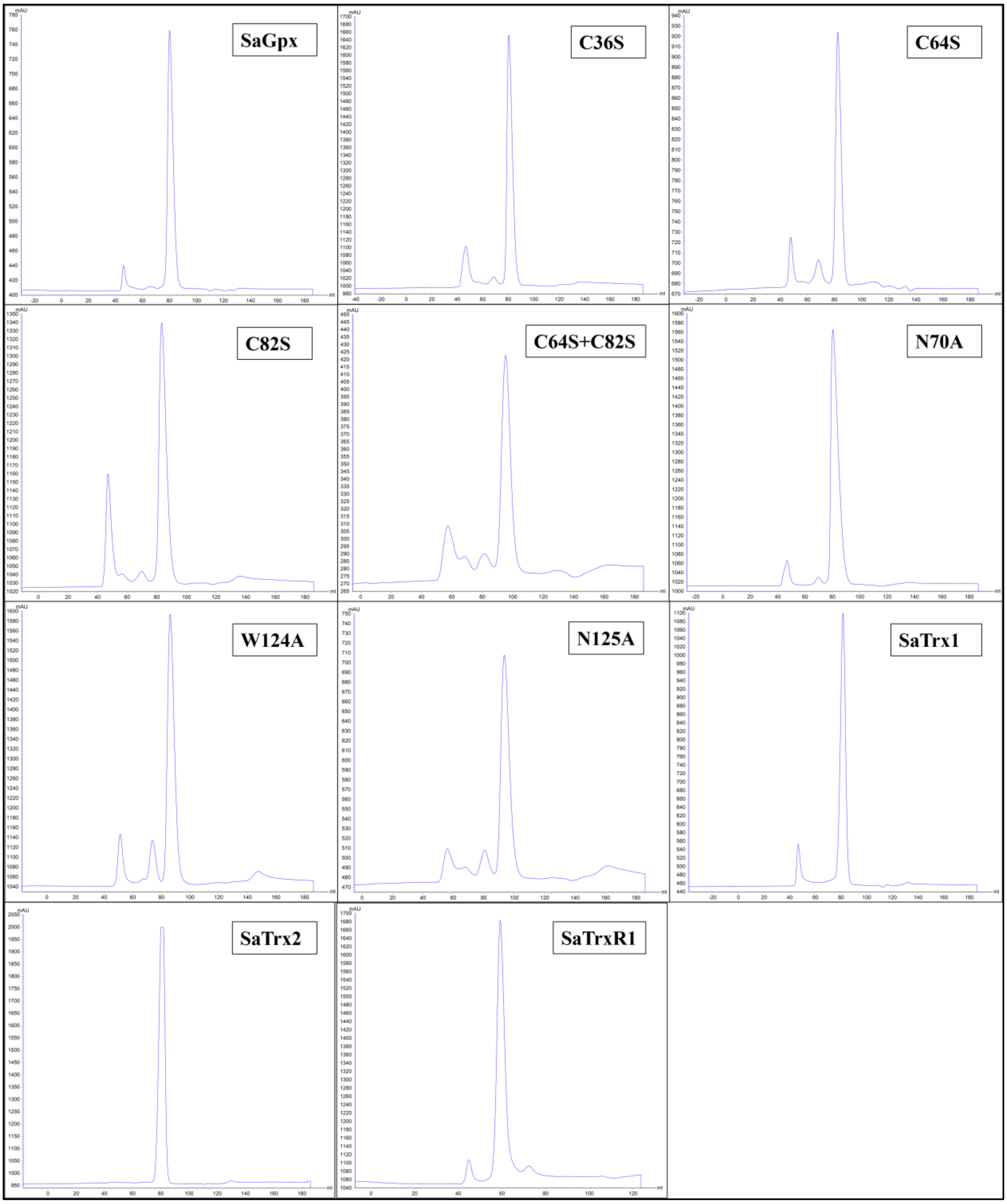


**Figure S1.** Size-exclusion chromatography profile of the homogenously purified proteins. Protein fraction showing single peak at $\lambda_{Max}$ = 280nm was collected and used for further biochemical characterization as well as crystallization purposes.

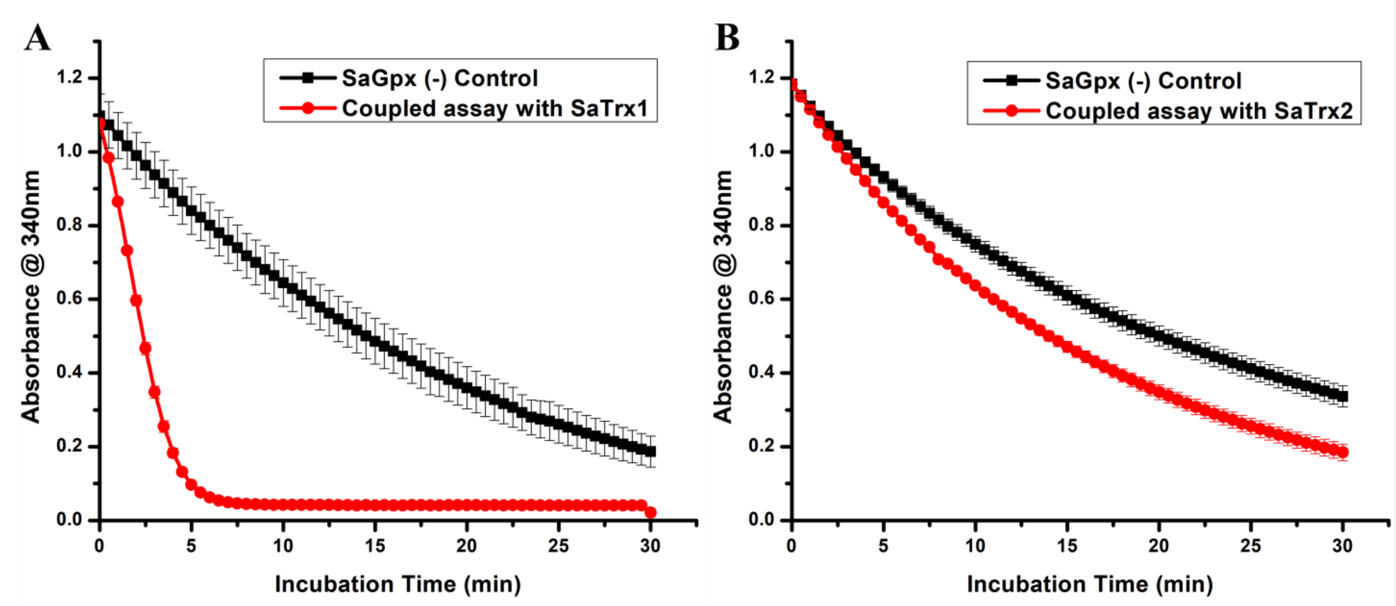


**Figure S2.** Time dependent coupled assay to probe cognate electron donor of SaGpx. Coupled assay perform with (A) SaTrx1 shows steep decrease in NADPH absorbance over time compared to (B) SaTrx2.

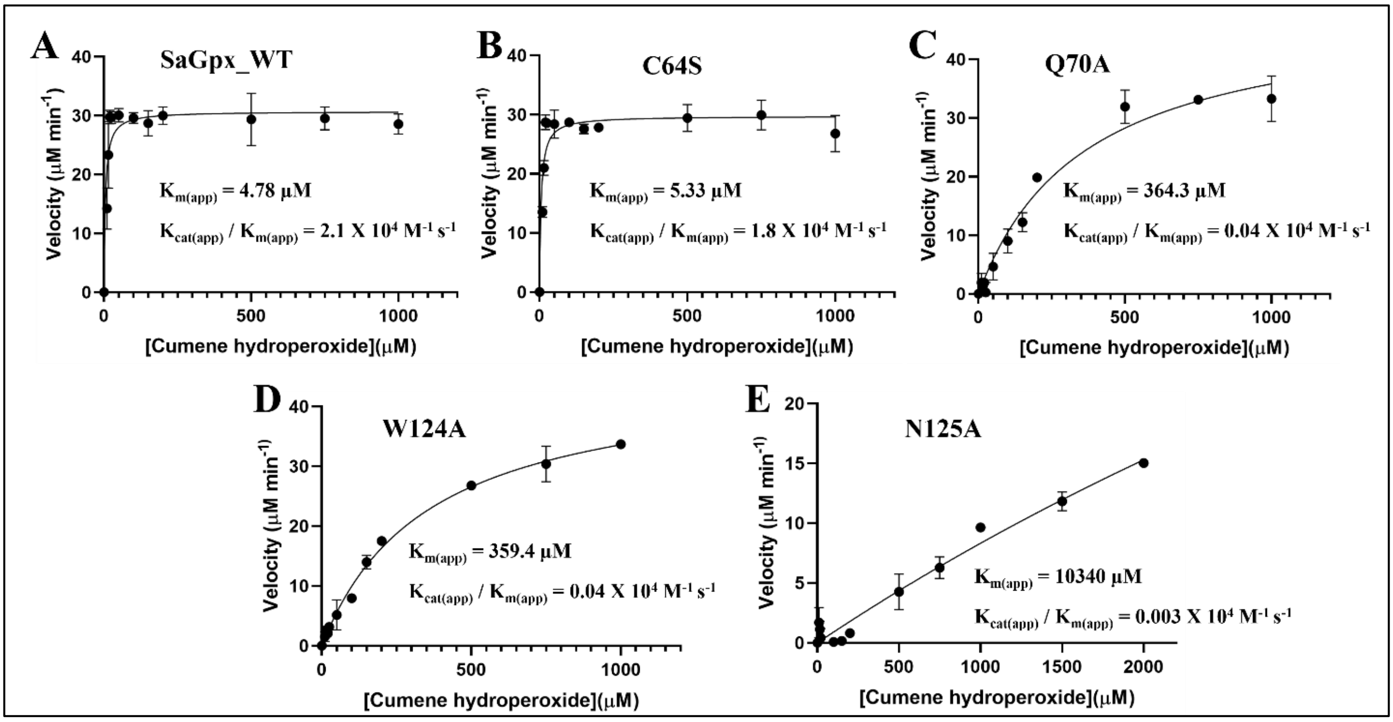


**Figure S3.** Michaelis-Menten kinetic characterization of different point mutants of SaGpx: (A) wild type SaGpx, (B) C64S SaGpx, (C) Q70A SaGpx, (D) W124A SaGpx, and (E) N125A SaGpx. Reaction velocity was plotted against substrate concentration and fitted into Michaelis-Menten equation [$V=V_{max}*S/(k_m+S)$]. Different kinetic values are listed in Table 3.

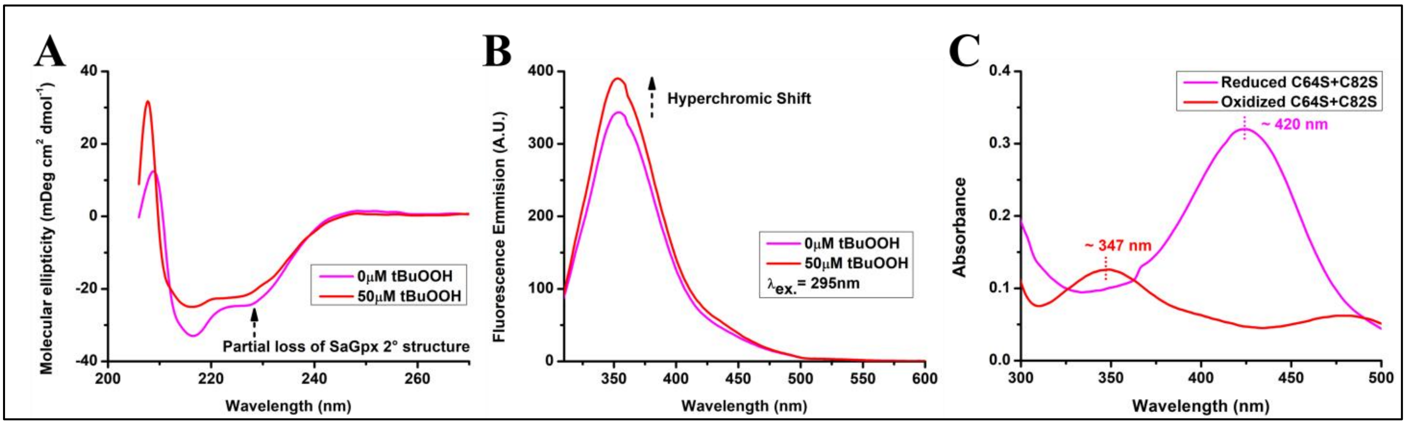


**Figure S4.** Spectroscopic analysis of SaGpx structural features. (A) Structural transition of SaGpx upon oxidation. CD spectra of SaGpx without (Magenta) and with t-BOOH (red). At 50μM t-BOOH concentration there is partial loss of secondary conformations which is reflected by the change in the characteristic ~222nm which in turn supports the structural change between the two redox states of SaGpx. (B) Fluorescence-based assessment of active site dynamics in SaGpx. Tryptophan fluorescence spectra of SaGpx exhibit a hyperchromic shift in the presence of t-BOOH (red), supporting conformational change in the proteins active site micro-environment. (C) Spectroscopic detection of sulfenic acid formation in SaGpx. The characteristic peak (at ~347 nm) for NBD-Cl adduct with sulfenic acid was observed in the oxidized protein sample (red) while no peak at ~420 nm. While the reduced protein sample shows only peak at ~420nm characteristic for NBD-Cl adduct with reduced thiol group. These collectively supports the formation of sulfenic acid intermediate during interaction with substrate hydroperoxide.

**Table S1.** Data collection and refinement statistics.

| Parameters | C36S mutant of SaGpx (PDB ID: 9KZM) |
|---|---|
| Wavelength | 0.978930 Å |
| Resolution range | 39.47 - 1.5 (1.55 - 1.5) |
| Space group | C 2 2 $2_1$ |
| Unit cell | 62.79 95.92 59.78 90 90 90 |
| Total reflections | 58536 (5739) |
| Unique reflections | 29273 (2871) |
| Multiplicity | 2.0 (2.0) |
| Completeness (%) | 99.86 (99.51) |
| Mean I/sigma(I) | 25.61 (3.38) |
| Wilson B-factor | 17.57 |
| R-merge | 0.01595 (0.2518) |
| R-meas | 0.02255 (0.3562) |
| R-pim | 0.01595 (0.2518) |
| CC1/2 | 1 (0.922) |
| CC* | 1 (0.98) |
| Reflections used in refinement | 29244 (2867) |
| Reflections used for R-free | 1476 (143) |
| R-work | 0.1505 (0.2368) |
| R-free | 0.1701 (0.2886) |
| CC(work) | 0.975 (0.949) |
| CC(free) | 0.969 (0.907) |
| Number of non-hydrogen atoms | 1473 |
| macromolecules | 1279 |
| ligands | 0 |
| solvent | 194 |
| Protein residues | 158 |
| RMS(bonds) | 0.019 |
| RMS(angles) | 1.51 |
| Ramachandran favored (%) | 98.72 |
| Ramachandran allowed (%) | 1.28 |
| Ramachandran outliers (%) | 0.00 |
| Rotamer outliers (%) | 0.00 |
| Clashscore | 1.97 |
| Average B-factor | 27.55 |
| macromolecules | 25.99 |
| solvent | 37.81 |
| Number of TLS groups | 1 |

Statistics for the highest-resolution shell are shown in parentheses.

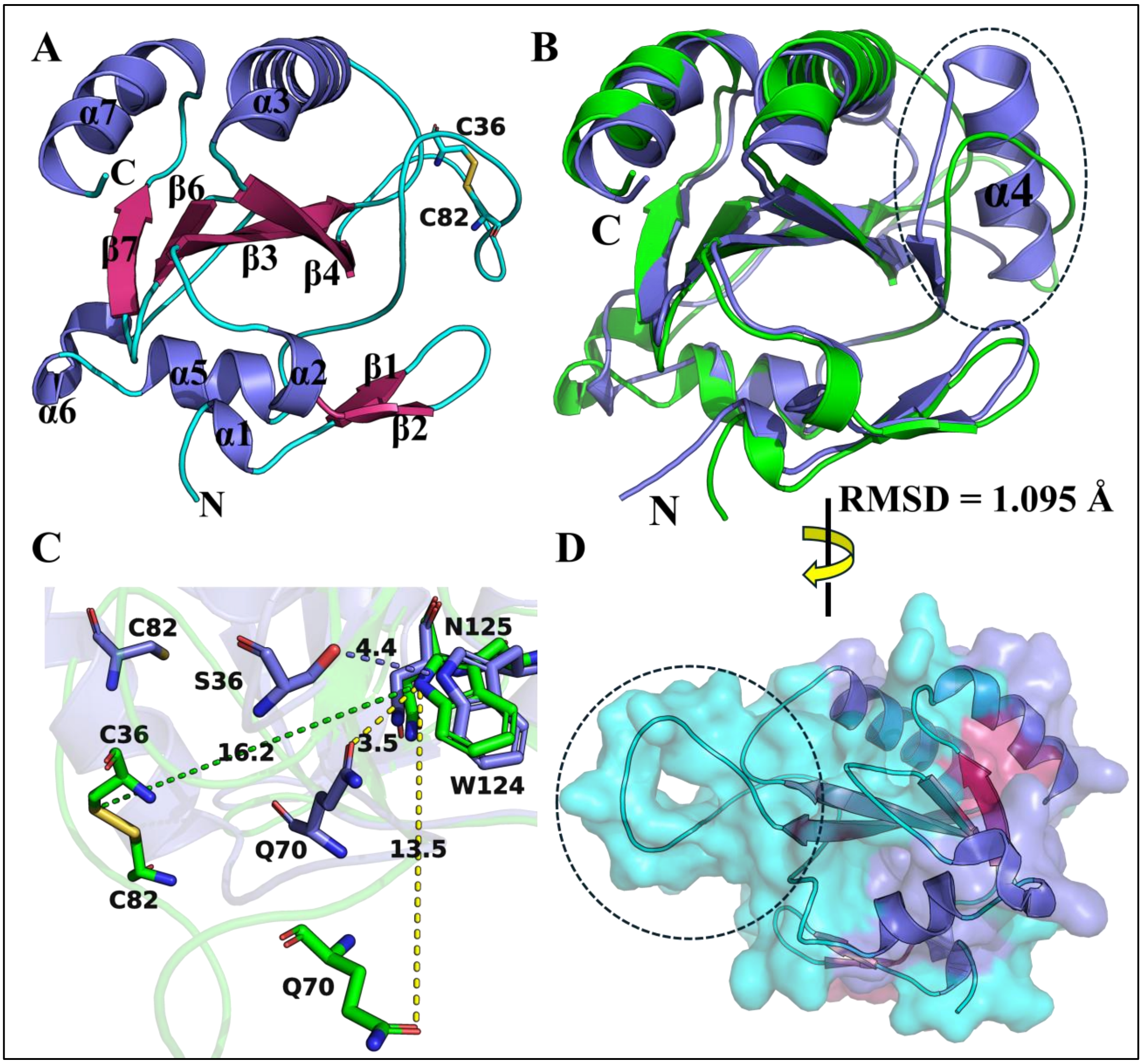


**Figure S5.** Model structure of Oxidized SaGpx obtained through homology modelling. Homology modelling was performed through Modeller software using oxidized crystal structure of poplar Gpx5 (PDB ID 2P5Q). (A) Cartoon representation of oxidized model of SaGpx. Model shows formation of disulfide bond between C36 and C82. (B) Structural superimposition of reduced crystal structure of SaGpx (C36S mutant) with modelled oxidized structure. Superimposition reveals the complete unwinding (highlighted by dotted black circle) of α-helix 4 (present in reduced state) into loop in oxidized state. This structural transition led to an RMSD value of 1.095Å. (C) Change in spatial arrangement of active site residues in both reduced and oxidized state of SaGpx. In oxidized state, due to structural transition, the polar residues like C36 and Q70 got shifted away from W124, leading to decrease in fluorescence quenching compared to reduced state. (D) Oxidation-induced change in surface feature of SaGpx. Conformational change in oxidized state generate surface features (highlighted by dotted black circle) that may be crucial in designing new enzyme inhibitors.

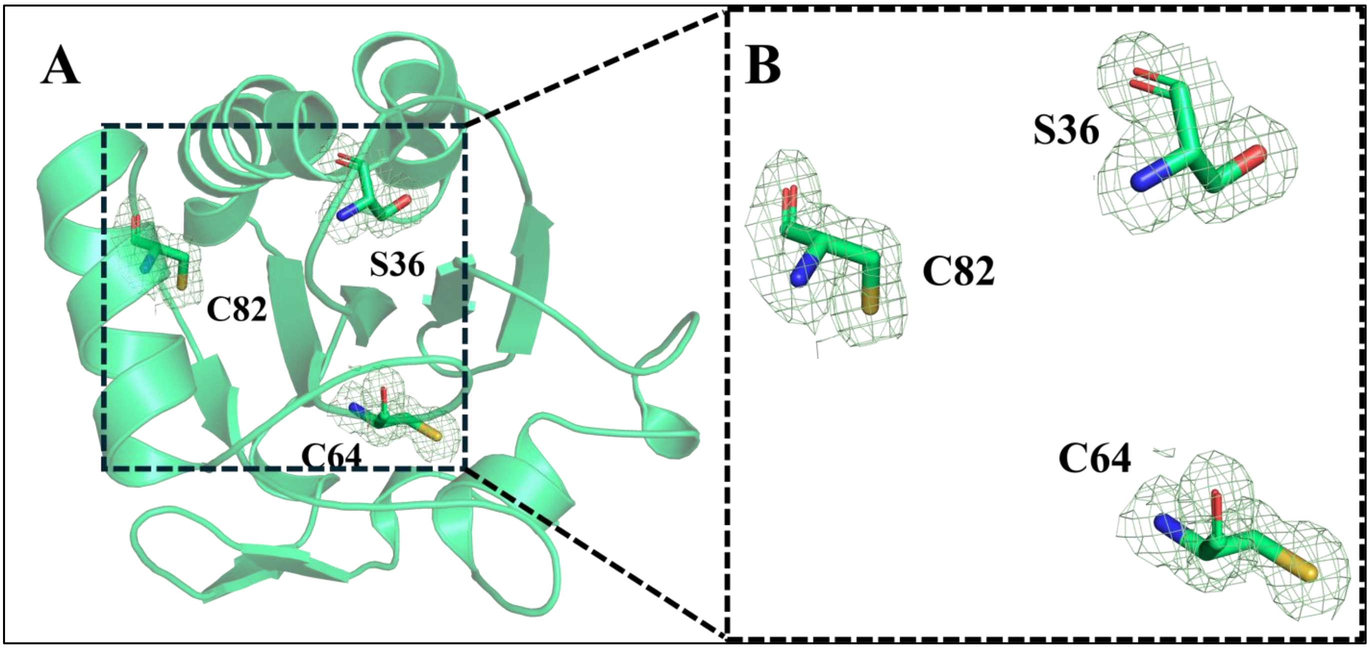


**Figure S6.** Structural analysis of putative resolving cysteine residues in the SaGpx C36S mutant crystal structure. (A) Overall spatial arrangement of C64 and C82 in the C36S mutant structure. (B) Close-up view showing electron density validation of the cysteine residues. The 2Fo-Fc electron density map (contoured at 1σ) clearly defines the sulfhydryl side chains of C64 and C82, confirming their presence in the reduced state.

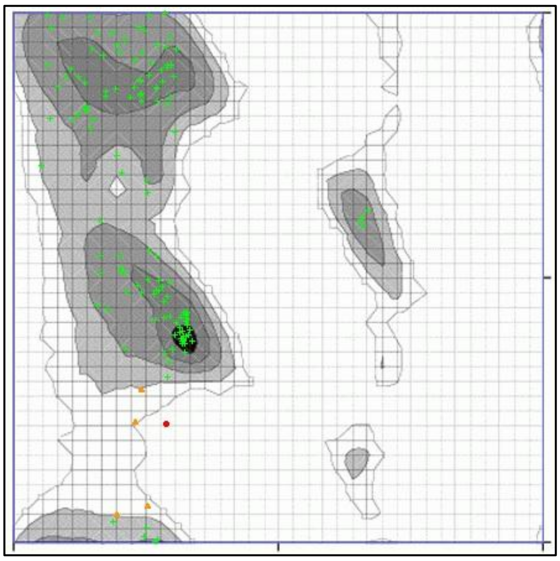

**Figure S7.** Ramachandran plot of oxidized model of SaGpx. Presence of all the amino acids in the allowed region supports the good quality of the model.

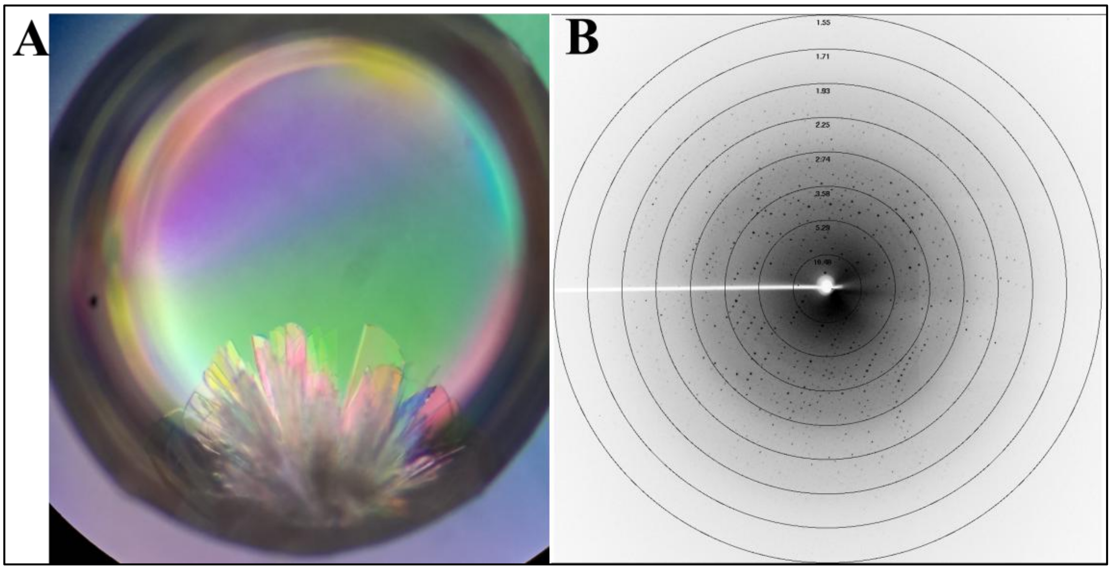


**Figure S8.** Crystals and single-crystal x-ray diffraction of SaGpx (C36S mutant). (A) Crystal drop shows plate like crystals radiating from a hub. Crystals obtained from hanging drop vapour diffusion method method. (B) X-ray diffraction pattern of single crystal. The well-defined sharp diffraction spots with low background noise indicate high crystal quality, with diffraction extending upto1.5Å.

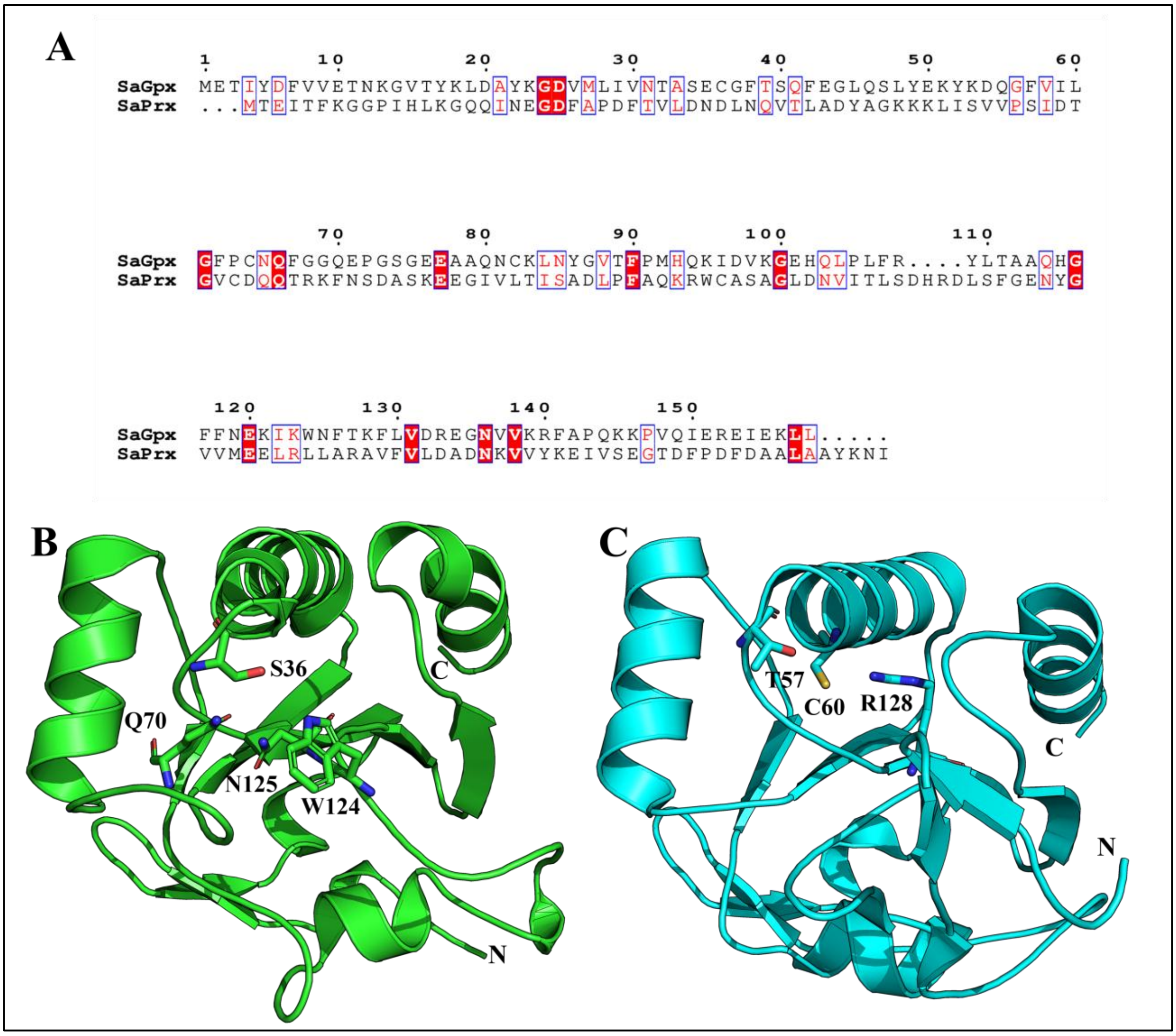


**Figure S9.** Comparison of SaGpx with Staphylococcal atypical 2-cys peroxiredoxin. (A) Multple sequence alignment of both the sequence. Cartoon representation of (B) SaGpx (green) and (C) SaPrx, Staphylococcal atypical 2-cys peroxiredoxin (cyan) (PDB Id: 3P7X) with an RMSD of 3.76 Å. Both sequential and structural analysis reveals very less similarity between these two proteins, however they share the same catalytic mechanism.
